# Type 2 conventional dendritic cells and regulatory T cells form a barrier tissue circuit to control allergic inflammation

**DOI:** 10.64898/2026.05.04.722795

**Authors:** Jenny M. Mannion, Lukas M. Altenburger, Hannah Willit, Neal P. Smith, Creel Ng Cashin, Jonah Clegg, Nandini Samanta, Victor Barrera, William J. Gammerdinger, Shannan Ho Sui, Caroline L. Sokol, Alexanda-Chloé Villani, Thorsten R. Mempel, Rod A. Rahimi

## Abstract

Chronic allergic diseases are driven by T helper type 2 (Th2) cells in barrier tissues. Despite their profound effects on tissue physiology, Th2 cells represent a rare cell population within tissues, suggesting mechanisms restraining Th2 cell expansion at barrier sites that remain ill defined. Using a murine model of allergic asthma, we demonstrate that effector Th2 cells promote cDC2 activation within the lungs, including expression of the CCR4 ligands that attract Foxp3^+^ regulatory T cells (Tregs). Selective deletion of *Ccr4* in Tregs during the effector Th2 cell response led to increased lung Th2 cells, activated cDC2s, and allergic inflammation.

Mechanistically, CCR4 promoted Treg trafficking efficiency and was required to specifically control tissue cDC2 co-stimulatory molecule expression. Lastly, in the airways of humans with allergy, the expression of the CCR4 ligands in activated cDCs correlated with Treg enrichment. In sum, we define a cDC2-Treg feedback circuit within a barrier tissue that restrains effector Th2 cell expansion, revealing a novel role for tissue cDC2s in controlling Th2 cell biology.

## Introduction

Type 2 immunity provides host protection against macroparasites, venoms, and toxins, but maladaptive type 2 immunity targeting noxious, environmental antigens causes allergic diseases.^1,2^ Specifically, allergic diseases are driven by the development of a population of allergen-specific, CD4^+^ T helper type 2 (Th2) cells with unique phenotypic and functional properties, which have been termed pathogenic effector Th2 cells or Th2A cells.^3–5^ In contrast to “conventional” Th2 cells, pathogenic Th2 cells are terminally differentiated and express higher levels of the type 2 cytokines IL-5, IL-9, and IL-13 that orchestrate allergic inflammation in barrier tissues.^4–7^ Notably, in humans, pathogenic Th2 cells are numerically rare at sites of allergic disease, suggesting mechanisms that limit the expansion of Th2 cells at barrier sites.^7–13^ Defining the endogenous mechanisms restraining pathogenic Th2 cells in barrier tissues has the potential to nominate novel therapeutic approaches for chronic allergic diseases.

The development of pathogenic Th2 cells requires instructive signals in the draining lymph node and barrier tissue. During priming in lymph nodes, a population of type 2 conventional dendritic cells (cDC2s), which are dependent on the transcription factors IRF4 and KLF4, initiate Th2 cell differentiation.^14–17^ Primed Th2 cells acquire the capacity to produce the type 2 cytokine IL-4, but do not exhibit significant expression of IL-5, IL-9, or IL-13.^18–20^ After primed Th2 cells egress from the draining lymph node, they traffic into barrier tissues where additional signals promote their differentiation into multi-cytokine producing Th2 cells.^21^ In the lungs, primed Th2 cells require direct stimulation by damage-associated molecular patterns (DAMPs) IL-33, TSLP, and IL-25 to differentiate into multi-cytokine producing effector cells.^20^ In addition, barrier tissue Th2 cells form clusters with MHCII^+^CD11c^+^ antigen-presenting cells (APCs) with depletion of CD11c^+^ APCs reducing the effector Th2 cell response, suggesting APC-Th2 cell clusters are required for effector Th2 cell differentiation and/or function.^22,23^ The tissue APC subsets regulating effector Th2 cells in vivo remain unclear. In mice, initial work suggested that monocyte-derived cells (MCs) in the lungs promoted effector Th2 cell responses.^24^ Subsequent single cell RNA-sequencing (scRNA-seq) approaches and flow cytometric analysis revealed that cDC2s in inflamed tissue had been mischaracterized as MCs due to shared phenotypic features.^25^ Furthermore, studies characterizing the barrier immune response in humans with allergic diseases have shown the enrichment of pathogenic Th2 cells and activated cDCs compared to controls.^7–9^ However, the role of cDCs versus MCs in controlling pathogenic Th2 cells in barrier tissues remains unclear.

Along with the enrichment of pathogenic Th2 cells and activated cDCs in barrier tissues during allergic inflammation, Foxp3^+^ regulatory T cells (Tregs) are also enriched at the sites of allergic disease in humans compared to controls.^7,10^ Foxp3^+^ Tregs are a unique population of CD4^+^ T cells that play an indispensable role in suppressing aberrant immune responses to self and environmental antigens, including allergens.^26–30^ Despite their importance in suppressing allergic immunity, the mechanisms whereby Tregs control Th2 cell responses in barrier tissues remain unclear. Tregs employ a range of suppressive mechanisms to inhibit APCs and effector T cells, including secretion of immunosuppressive cytokines such as IL-10, IL-35, and TGF-β, sequestration of IL-2, blocking or stripping co-stimulatory molecules and peptide:MHCII complexes from APCs, among others.^31^ For Tregs to execute targeted suppressive function, they must localize to the sites of APC-T cell crosstalk.^32^ The activated cDCs enriched in barrier tissues during allergic inflammation are characterized by high expression of the CCR4 ligands, *CCL17* and *CCL22*.^7–9^ The chemokine receptor CCR4 is expressed by both Th2 cells and Tregs in humans and mice with adoptive transfer studies suggesting CCR4 is required for Th2 cell and Treg trafficking into barrier tissues.^33–37^ As the CCR4 system co-localizes Th2 cells and Tregs, we posited that interrogating the CCR4 system would reveal novel biology regarding the mechanisms controlling pathogenic Th2 cells in vivo.

Here, we use a murine model of allergic lung inflammation to define tissue DC crosstalk between Th2 cells and Tregs in a barrier tissue. Using scRNA-seq and flow cytometric analysis, we demonstrate that allergic inflammation leads to a dramatic expansion of activated cDC2s expressing PD-L2 and CCR7 within the lungs, which are the dominant DC source of the CCR4 ligands during allergic inflammation. We show that the innate immune response to allergen was insufficient to promote the expansion of activated cDC2s within the lungs. Rather, CD4^+^ T cells and IL-4 and IL-13 were required to promote cDC2 activation and CCR4 ligand expression, suggesting a feedforward circuit between effector Th2 cells and cDC2s. Inducible and selective deletion of *Ccr4* in Tregs during allergic inflammation led to a dramatic increase in lung Th2 cells, cDC2s, and allergic inflammation without any significant effect in the draining lymph node. Notably, CCR4 was not absolutely required for Treg transmigration into the lungs or positioning within cDC2-CD4^+^ T cell clusters during allergic inflammation, but was required to restrain the expansion of cDC2-CD4^+^ T cell clusters in the lungs. Mixed bone marrow chimera experiments revealed that deletion of CCR4 led to reduced Treg trafficking efficiency. In support of a model in which CCR4 specifically controls Treg trafficking efficiency, RNA-seq analysis demonstrated that deletion of *Ccr4* in Tregs did not affect the core Treg program or immunosuppressive cytokine expression, but led to reduced expression of *Asb2*, an E3 ubiquitin ligase that promotes integrin-dependent cell trafficking.^38^ We demonstrate that IL-4 directly induces *Asb2* expression in Tregs and IL-4 signaling promotes Treg trafficking efficiency in a cell-intrinsic manner, suggesting that CCR4 and IL-4 collaborate to promote Treg trafficking to the sites of allergic immunity. In addition, we show that *Ccr4* deletion in Tregs leads to enhanced expression of CD80 and CD86 in tissue cDC2s, but not MCs, demonstrating that Tregs require CCR4 to specifically control co-stimulatory molecule expression in tissue cDC2s. Lastly, to support our findings in humans, we analyzed scRNA-seq data of lower airway mucosal cells in a human model of allergen-induced inflammation, demonstrating that the expression of the CCR4 ligands in activated cDCs correlated with Treg enrichment. In sum, we define a cDC2-Treg feedback circuit within a barrier tissue that controls effector Th2 cells and allergic inflammation, unveiling a novel role for tissue cDC2s in controlling the expansion of Th2 cells and the severity of allergic disease.

## Results

### Activated cDC2s are the dominant lung DC subset expressing *Ccl17* and *Ccl22* during allergic inflammation

To characterize the lung DCs expressing the CCR4 ligands, we used a well-established, murine model of allergic asthma via intranasal house dust mite extract (HDM) administration (day 0 sensitization, days 7-11 challenges) (Figure 1A).^19,24^ On day 14, which represents peak allergic inflammation, we confirmed that HDM induces expression of the CCR4 ligands within the lungs by performing RT-qPCR on total lung RNA for *Ccl17* and *Ccl22*. Consistent with published studies, we found allergic inflammation promotes a significant increase in both CCR4 ligands (Figure S1A).^39–41^ Given that DCs are the dominant source of the CCR4 ligands during allergic inflammation in mice and humans, we sought to determine the lung DC subsets expressing *Ccl17* and *Ccl22*.^7–9,40^ First, we quantitated the lung DC subsets during peak allergic inflammation by flow cytometry. On day 14 of the HDM model, we performed i.v. injection with fluorophore-labeled anti-CD45 antibody to label intravascular leukocytes before lung harvest and analyzed intraparenchymal lung CD11c^+^MHCII^+^ cells via flow cytometry.^8^ We used the marker XCR1 to identify cDC1s as well as CD26 and CD64 to distinguish cDC2s and MCs, respectively (Figure S1B).^25^ Compared to naïve mice, HDM-treated mice exhibited a significant expansion of lung cDC2s and MCs without any increase in cDC1s (Figure 1B). Next, to determine which of these lung DC subsets express the CCR4 ligands, we performed scRNA-seq analysis of lung intraparenchymal CD11c^+^MHCII^+^ cells sorted from naïve and HDM-treated mice on day 14. Analysis of 23,592 total DCs from naïve and HDM-treated mice identified 5 clusters of cDCs and 3 clusters of MCs (Figure 1C). Specifically, we identified clusters of “cycling” cDC1s and cDC2s that expressed cell cycle genes and likely represent cells that differentiated from recently recruited and proliferating pre-cDCs as previously described.^42^ We also identified cDC1s and cDC2s lacking expression of cell cycle genes as well as activated/mature *Ccr7*-expressing cDCs. Among the MC clusters, we identified a large cluster of cells with DC features without canonical macrophage markers that we annotated as monocyte-derived DCs (MoDCs) as well as macrophages and a small cluster of cells expressing alveolar macrophage markers and likely representing monocyte-derived alveolar macrophages that increase during lung inflammation (Figure 1C). Consistent with our flow cytometry data (Figure 1B), compared to naïve mice, HDM-treated mice displayed an enrichment of the cDC2 and MC clusters with decreased enrichment in the cDC1 cluster (Figure 1C). Next, we evaluated the expression of *Ccl17* and *Ccl22* in the clusters in naive and HDM-treated mice. Naïve mice exhibited basal expression of *Ccl17* predominantly in cDC1s, cDC2s, and *Ccr7* cDCs whereas *Ccl22* exhibited restricted expression to *Ccr7* cDCs (Figure 1D, S1C and S1D). In HDM-treated mice, there was a notable increase in *Ccl17* expression across the cDC2s and *Ccr7* cDC clusters with minimal expression changes in cDC1s and MC clusters compared to naive mice (Figure 1D, S1C and S1D). In contrast, *Ccl22* expression remained restricted to activated *Ccr7* cDCs (Figures 1D, S1C, and S1D). Overall, *Ccr7*-expressing cDCs expressed the highest levels of *Ccl17* and *Ccl22* on a per cell basis in naïve mice, which increased further during allergic inflammation (Figures 1D, S1C and S1D). Given that cDCs downregulate subset-defining transcripts during activation, we were unable to determine the extent to which *Ccr7*-expressing cDCs were derived from cDC1s or cDC2s in our scRNA-seq dataset.^43^ As activated cDCs maintain surface protein expression of subset-defining markers, we used flow cytometry to define the extent to which CCR7^+^ cDCs exhibit a cDC1 or cDC2 phenotype during allergic inflammation. We found that virtually all of the CCR7^+^ cDCs in the lungs during allergic inflammation are cDC2s, which express higher levels of the DC activation marker PD-L2 than other DC subsets (Figures 1E-F). Consequently, our results suggest that activated, PD-L2^+^CCR7^+^ cDC2s are the dominant lung DC subset promoting chemotaxis of CCR4-expressing T cells during allergic inflammation.

**Figure 1.**
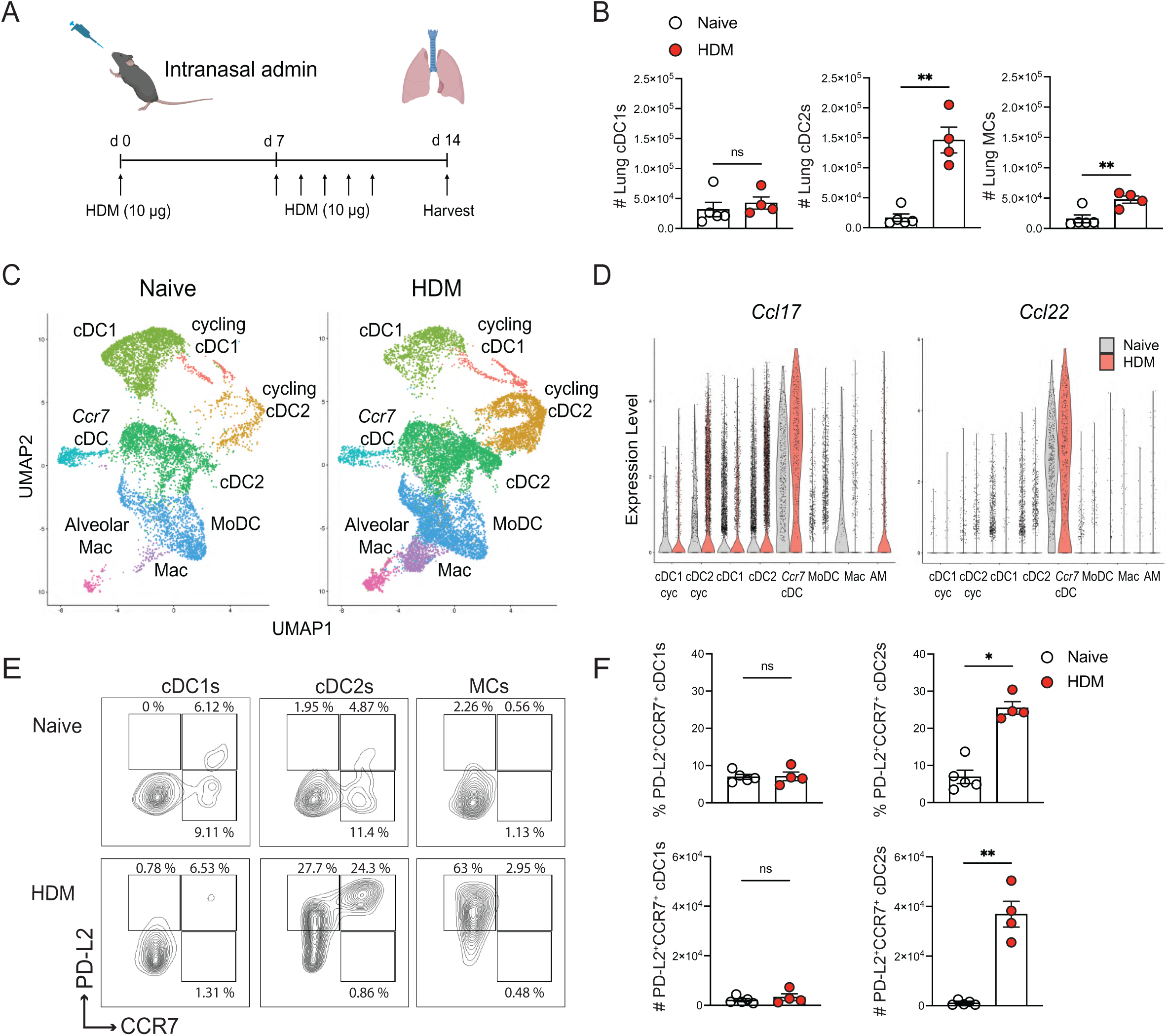
Lung CCR7+ cDC2s are the dominant DC subset expressing *Ccl17* and *Ccl22* during allergic inflammation. C57BL/6 mice were sensitized (day 0) with 10 µg i.n. HDM followed by daily challenges of 10 µg i.n. HDM on days 7–11 then injected with anti-CD45 antibody i.v. 3 minutes prior to tissue harvest on day 14. **A.** Acute intranasal HDM treatment protocol. **B.** Lung intraparenchymal (anti-CD45 i.v. antibody negative) DC subsets from naïve and HDM-treated mice, quantified via flow cytometry. **C.** scRNA-seq UMAPs with annotated lung DC clusters from naïve and HDM-treated mice. **D.** Violin plots showing expression of *Ccl17* and *Ccl22* in lung DC subsets from naïve and HDM-treated mice. **E.** Representative flow cytometry of lung DCs showing PD-L2 and CCR7 expression from indicated groups. **F**. Quantification of lung CCR7^+^PD-L2^+^ cDC1s and cDC2s from indicated groups. Representative data shows individual mice with mean ± SEM from one of three independent experiments with *n*L=L5 mice per group (B & F) or pooled from two independent experiments, *n* = 10 mice per group (C & D). For statistical analysis, a two-tailed *t* test was performed for parametric data, and a two-tailed Mann-Whitney *U* test was performed for nonparametric data. *, p<0.05; **, p<0.01; ***, p<0.001; ns, not significant.

### CD4^+^ T cells and IL-4Rα-dependent signaling are required for lung cDC2 activation during allergic inflammation

Given that allergic inflammation promotes the expansion of PD-L2^+^CCR7^+^ cDC2s expressing the CCR4 ligands (Figure 1), we sought to determine how allergen treatment promotes lung cDC2 activation. Previous studies investigating the role of allergen-induced DC activation have focused on initial sensitization, demonstrating that allergens promote DC activation by inducing production of alarmins such as TSLP and IL-33, which either directly act on DCs or promote ILC2 production of IL-13 that induces DC expression of CCL17.^39,44–49^ To characterize the role of the innate versus adaptive response to HDM in promoting cDC2 activation at the peak of allergic inflammation, we used C57BL/6 and MHCII knockout mice, which both possess alarmins and ILC2s, but the latter lacks CD4^+^ T cells. We treated C57BL/6 (control, ctrl) and MHCII knockout mice with intranasal HDM (days 0, 7-11). On day 14, we harvested lungs and performed flow cytometry on total CD11c^+^ cells to quantitate the intraparenchymal lung DC subsets. Consistent with published work, we found that MHCII-deficient mice exhibited significantly fewer lung eosinophils compared to controls (Figure S2A).^50^ Compared to controls, MHCII-deficient mice exhibited reduced total cDC2s and a dramatic reduction in PD-L2^+^CCR7^+^ cDC2s in the lungs on day 14 (Figure 2A-B). We did not detect any significant decrease in cDC1 or MC populations in mice lacking MHCII when compared to controls (Figure S2A). To identify pathways whereby the adaptive immune response promotes cDC2 activation, we analyzed our lung DC scRNA-seq dataset, assessing the expression of type 2 cytokine receptors that could explain our findings that CD4^+^ T cells promote cDC2, but not cDC1, activation. We found that *Il4r*α expression, which encodes the shared component of the IL-4 and IL-13 receptors, was expressed in cDC2s but not cDC1s during allergic inflammation (Figure 2C). Next, we asked whether IL-4R signaling is sufficient to promote DC activation. We cultured bone marrow-derived dendritic cells (BMDCs) in the presence and absence of recombinant IL-4 and measured expression of PD-L2 and CCR7. We found that IL-4 promoted a significant increase in the frequency of PD-L2^+^CCR7^+^ DCs in BMDCs treated with IL-4 when compared to controls, indicating that IL-4 is sufficient to promote DC activation in vitro (Figures 2D-E). To determine whether cDC2 expansion and activation are dependent on signaling via IL-4Rα in vivo, we treated control and *Il4r*α knockout mice with intranasal HDM (days 0, 7-11). On day 14, we harvested lungs and performed flow cytometry to quantitate the lung DC subsets. As expected, we found that mice deficient in IL-4Rα displayed significantly fewer lung eosinophils than control mice (Figure S2B).^51–53^ Notably, mice lacking IL-4Rα displayed a dramatic reduction in the accumulation of lung cDC2s as well as activated PD-L2^+^CCR7^+^ cDC2s without any influence on lung cDC1 or MC numbers (Figures 2F-G; S2B). Given that *Il4ra* knockout mice are broadly defective in the development of an adaptive type 2 immune response, we next asked whether specifically blocking IL-4 and IL-13 during the allergic response would impair cDC2 accumulation and activation in the lungs. We treated C57BL/6 mice with HDM (days 0, 7-11) and either control IgG or anti-IL-4/anti-IL-13 antibodies via intraperitoneal injection late in the HDM challenge phase on days 9, 11, 13. We found that blocking IL-4 and IL-13 resulted in a significant reduction in lung eosinophils, confirming effective blockade of IL-4 and IL-13 (Figure S2C). While the expansion of lung cDC2s was similar between the groups, blocking IL-4 and IL-13 led to a significant reduction in PD-L2^+^CCR7^+^ cDC2s (Figure 2H-I). Unexpectedly, the blockade of IL-4 and IL-13 led to an increase in the number of lung cDC1s, suggesting that the type 2 cytokines restrain the expansion of cDC1s during allergic inflammation (Figure S2C).

**Figure 2.**
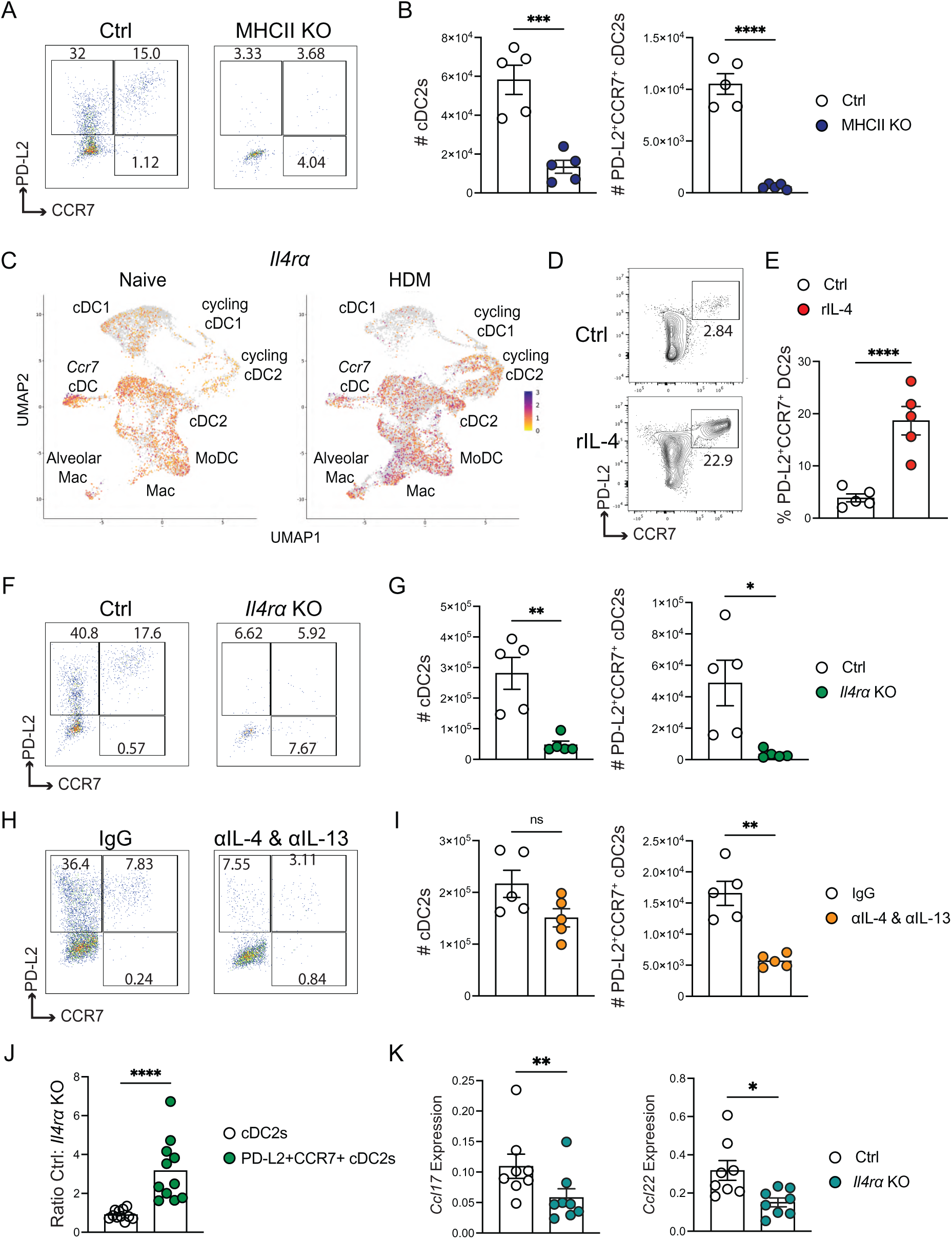
CD4+ T cells and IL-4/IL-13 signaling are required for lung cDC2 activation during allergic inflammation. Indicated mice were sensitized (day 0) with 10 µg i.n. HDM followed by daily challenges of 10 µg i.n. HDM on days 7–11 then injected with anti-CD45 antibody i.v. 3 minutes prior to tissue harvest on day 14. **A.** Representative flow cytometry of lung cDC2s from HDM-treated C57BL/6 control (Ctrl) and MHCII KO mice demonstrating PD-L2 and CCR7 expression. **B.** Number of total lung cDC2s and PD-L2^+^CCR7^+^ cDC2s from HDM-treated Ctrl and MHCII KO mice. **C.** UMAP of *Il4ra* expression in lung DC clusters from nailJve and HDM-treated mice. **D-E**. Bone marrow-derived cells were cultured for 6 days in media containing GM-CSF to generate bone marrow-derived dendritic cells (BMDCs) followed by stimulation with vehicle (Ctrl) or rIL-4 20 ng/ml for 4 days. **D.** Representative flow cytometry of BMDCs demonstrating PD-L2 and CCR7 expression. **E.** Frequency of PD-L2^+^CCR7^+^ cells among CD11c^+^ BMDCs from indicated groups. **F.** Representative flow cytometry of lung cDC2s from HDM-treated Ctrl and *Il4ra* KO mice demonstrating PD-L2 and CCR7 expression. **G**. Number of total lung cDC2s and PD-L2^+^CCR7^+^ cDC2s from HDM-treated Ctrl and *Il4ra* KO mice. **H.** Representative flow cytometry of lung cDC2s from mice treated with HDM and IgG or anti-IL-4/IL-13 antibodies via intraperitoneal (i.p) injection on days 7,11, and 13 of HDM treatment protocol. **I.** Number of total lung cDC2s and PD-L2^+^CCR7^+^ cDC2s from HDM-treated mice administered IgG or anti-IL-4/IL-13 antibodies **J-K**. Mixed bone marrow chimeras were generated with Ctrl and *Il4r*α-deficient bone marrow cells. Ctrl:*Il4ra*-deficient mixed bone marrow chimeras were treated with HDM (days 0, 7-11), followed by anti-CD45 antibody i.v. and tissue harvest on day 14. **J**. Ctrl:*Il4ra* KO ratio of indicated lung cDC2 subsets from mixed bone marrow chimera mice. **K.** *Ccl17* and *Ccl22* relative RNA expression in sorted lung cDC2s from Ctrl and *Il4ra* KO donors in mixed bone marrow chimera mice. Representative data shows individual mice with mean ± SEM from one of 2-3 independent experiments with *n*L=L5 mice per group, 5 independent experiments (E), or pooled from two independent experiments, *n* = 11 (J) or 8 (K) mice. For statistical analysis, a two-tailed *t* test was performed for parametric data, and a two-tailed Mann-Whitney *U* test was performed for nonparametric data. *, p<0.05; **, p<0.01; ***, p<0.001; ****, p<0.0001; ns, not significant.

Since cDC1s exhibit low expression of *Il4ra*, the inhibitory mechanism is likely cell-extrinsic. To determine the extent to which IL-4/IL-13 signaling promotes cDC2 activation in a cell-intrinsic manner, we generated mixed bone marrow chimeras with a 4:1 mix of control and *Il4r*α knockout (KO) bone marrow cells (Figure S2D). We treated control:*Il4r*α KO mixed bone marrow chimeras with HDM (days 0, 7-11). On day 14, we performed flow cytometry for lung DC subsets and found no difference in the ratio (normalized to total splenic CD45^+^ cells) of control:*Il4ra* KO cDC2s, indicating IL-4/IL-13 signaling does not directly regulate lung cDC2 subset expansion during allergic inflammation (Figures 2J). However, we observed a significant increase in the ratio of control:*Il4r*α KO cells among PD-L2^+^CCR7^+^ cDC2s (Figure 2J), demonstrating that IL-4/IL-13 signaling directly promotes cDC2 activation in a cell-intrinsic manner. IL4Rα-dependent signaling did not influence the accumulation of lung cDC1s or MCs (Figure S2E). To determine the requirement for *Il4r*α signaling in the expression of the CCR4 ligands, we used congenic markers to sort CD45.1^+^ control and CD45.2^+^ *Il4r*α KO cDC2s from control:*Il4ra* KO chimeric mice and analyzed *Ccl17* and *Ccl22* expression via RT-qPCR (Figure S2F). cDC2s lacking IL-4Rα displayed significantly reduced expression of the CCR4 ligands compared to control cDC2s from the same inflammatory environment (Figure 2K). Taken together, these data indicate that during the effector phase driving allergic inflammation, Th2 cell-derived cytokines IL-4 and IL-13 signal directly through IL-4Rα to promote cDC2 activation and CCR4 ligand production.

### Regulatory T cells require CCR4 to restrain lung Th2 cells, activated cDC2s, and allergic inflammation

Our results above indicate that effector Th2 cells drive tissue cDC2 activation, including expression of the CCR4 ligands. Given that effector Th2 cells are numerically rare at sites of allergic disease, we posited that the feedforward circuit between effector Th2 cells and cDC2s must be sensed and restrained. While CCR4 ligand expression during allergic inflammation is canonically viewed as a mechanism for Th2 cell recruitment, Tregs also express CCR4. We sought to determine the role of CCR4 in Treg control of activated cDC2s, Th2 cells, and allergic inflammation. First, we phenotyped CCR4-expressing Tregs in the lung parenchyma during allergic inflammation. On day 14 of the acute HDM model, we found that approximately 40% of lung intraparenchymal Tregs exhibited surface expression of CCR4 (Figure S3A-B). Compared to CCR4^-^ Tregs, CCR4^+^ Tregs expressed higher levels of the activation marker CD44 as well as numerous Treg effector and co-stimulatory molecules, including TIGIT, GITR, and ICOS (Figures S3C-D). These findings demonstrate that CCR4^+^ Tregs exhibit an activated phenotype, suggesting they are poised for executing suppressive mechanisms. We did not detect any significant difference in expression of the IL-33 receptor, ST2 in CCR4^+^ Tregs compared to CCR4^-^ Tregs (Figure S3D). We next sought to determine the role of CCR4 in Treg suppressive function in vivo. Selectively defining the role of CCR4 in Tregs during allergic inflammation has been challenging due to a lack of genetic tools for conditional *Ccr4* deletion.^34,36,37^ Specifically, global *Ccr4* knockout mice do not distinguish the role of CCR4 in Th2 cells versus Tregs.

Furthermore, global *Ccr4* knockout mice exhibit defects in central tolerance and develop spontaneous autoimmunity, limiting the use of this model to study the allergic response in vivo.^54,55^ To investigate the function of CCR4 expression specifically in Tregs in an inducible manner, we generated a novel *Ccr4* floxed (fl) mouse. We crossed our *Ccr4^fl^* mice to tamoxifen-inducible *Foxp3^EGFP-Cre-ERT2^*mice.^56^ To selectively delete *Ccr4* in Foxp3^+^ Tregs during the HDM challenge phase, we treated *Foxp3^EGFP-Cre-ERT2^* control mice and *Foxp3^Cre-ERT2^* x *Ccr4^fl/fl^* mice with HDM (days 0, 7-11) as well as tamoxifen daily via oral gavage from days 4-8 (Figure 3A).

**Figure 3.**
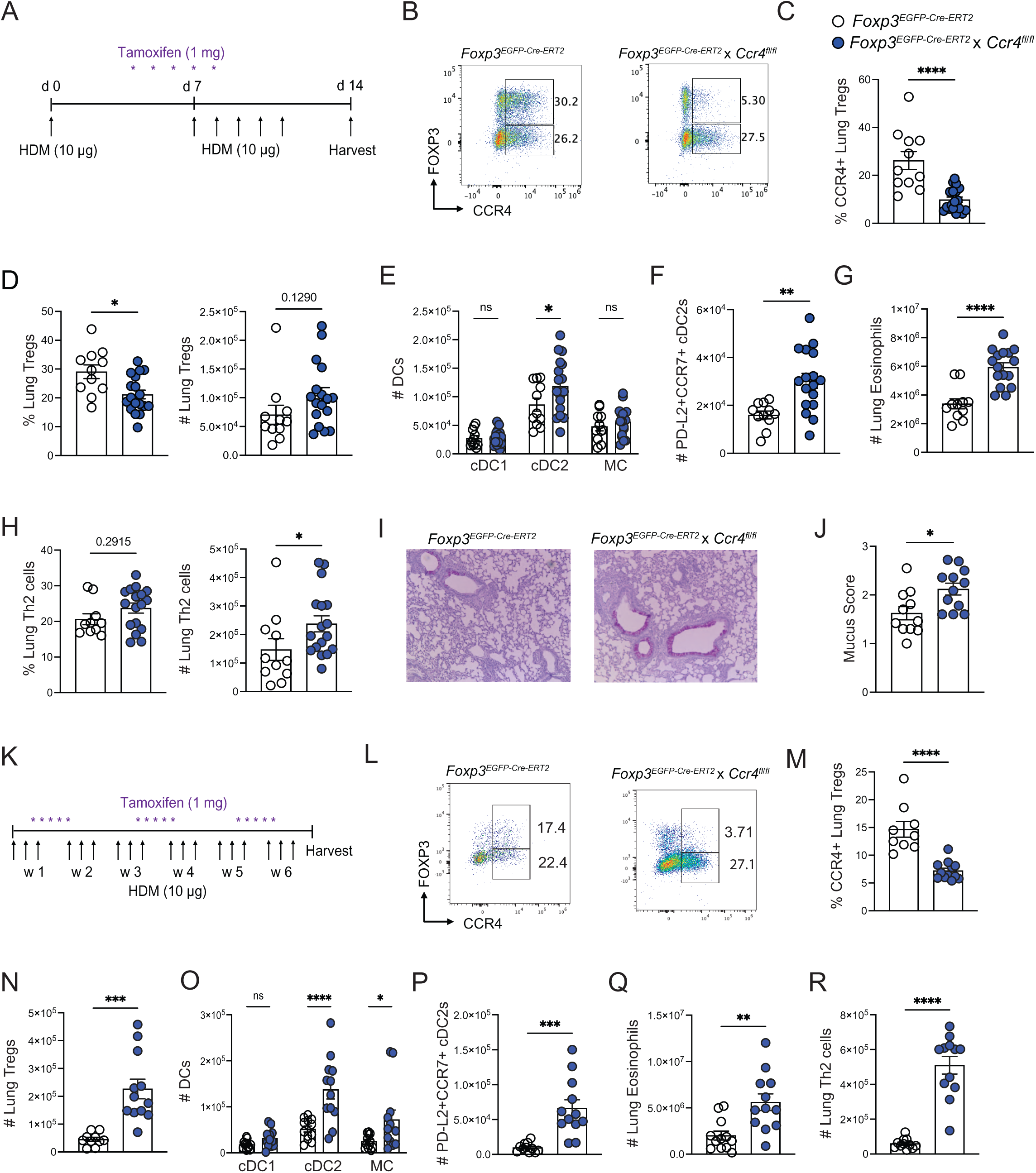
Regulatory T cells require CCR4 to restrain lung Th2 cells, activated cDC2s, and allergic inflammation. *Foxp3^EGFP-Cre-ERT2^* control mice and *Foxp3^EGFP-Cre-ERT2^* x *Ccr4^fl/fl^* mice were sensitized with 10 µg i.n. HDM followed by daily challenges of 10 µg i.n. HDM on days 7–11, 1 mg daily doses of tamoxifen via oral gavage on days 4-8, then injected with anti-CD45 antibody i.v. 3 minutes prior to tissue harvest on day 14. **A.** Acute intranasal HDM and tamoxifen treatment protocol. **B.** Representative flow cytometry of lung intraparenchymal (anti-CD45 i.v. antibody negative) CD4^+^ T cells showing Foxp3 and CCR4 expression from indicated groups. **C.** Percent lung CCR4^+^Foxp3^+^ Tregs among Foxp3^+^ Tregs from indicated groups. **D.** Percentage of Foxp3^+^ Tregs among lung CD4^+^ T cells and number of total lung Tregs from indicated groups. **E.** Number of total lung DC subsets from indicated groups. **F.** Number of PD-L2^+^CCR7^+^ lung cDC2s. **G.** Number of lung eosinophils **H.** Percentage of Foxp3^-^ST2^+^ T cells (Th2 cells) among lung CD4^+^ T cells and number of total lung Th2 cells**. I.** Representative PAS-stained lung sections. **J.** Mucus scores. *Foxp3^EGFP-Cre-ERT2^* control mice and *Foxp3^EGFP-Cre-ERT2^* x *Ccr4^fl/fl^* mice were administered 10 µg i.n HDM 3 times per week for 6 weeks alongside 1 mg daily doses of tamoxifen via oral gavage on days 4-8, 18-22 and 32-36 of HDM treatment then injected with anti-CD45 antibody i.v. 3 minutes prior to tissue harvest on day 42. **K.** Chronic intranasal HDM and tamoxifen treatment protocol. **L.** Representative flow cytometry of lung CD4^+^ T cells showing Foxp3 and CCR4 expression from indicated groups. **M.** Percentage of lung CCR4^+^Foxp3^+^ Tregs. **N.** Number of lung Tregs. **O.** Number of lung DC subsets. **P.** Number of PD-L2^+^CCR7^+^ lung cDC2s. **Q.** Number of lung eosinophils **R.** Number of lung Th2 cells. Data are from three independent experiments with 11-18 mice pooled (A-J) or data are from two independent experiments with 12 mice pooled (K-R). For statistical analysis, a two-tailed *t* test was performed for parametric data, and a two-tailed Mann-Whitney *U* test was performed for nonparametric data. One-way ANOVA analysis with Holm-Sidak’s testing for multiple comparisons. *, p<0.05; **, p<0.01; ***, p<0.001; ****, p<0.0001; ns, not significant.

Compared to controls, *Foxp3^EGFP-Cre-ERT2^* x *Ccr4^fl/fl^* mice exhibited selective deletion of CCR4 in Tregs without influencing CCR4 expression in conventional Foxp3^-^CD4^+^ T cells (Figures 3B-C). Unexpectedly, deletion of CCR4 in Tregs did not reduce the number of Tregs in the lung parenchyma, but rather there was a trend towards an increased number of total lung Tregs compared to controls (Figure 3D). Furthermore, we did not observe any significant difference in Treg expression of Ki67, TIGIT, GITR, ICOS or ST2 in *Foxp3^EGFP-Cre-ERT2^* x *Ccr4^fl/fl^* mice compared to controls (Figure S4A). These results suggest that CCR4 is not required for Treg recruitment, proliferative potential, or co-stimulatory/inhibitory molecule expression. In terms of the effect on Treg suppressive function within the lungs, compared to controls, deletion of *Ccr4* in Tregs led to a specific increase in the number of lung cDC2s with no difference in the number of cDC1s or MCs (Figure 3E). Additionally, we detected a significant increase in the percentage and number of activated PD-L2^+^CCR7^+^ cDC2s in *Foxp3^EGFP-Cre-ERT2^* x *Ccr4^fl/fl^* mice compared to controls (Figures S4B and 3F). We next wanted to assess the extent to which CCR4 expression in Tregs is required to control allergic inflammation. Compared to controls, *Foxp3^EGFP-Cre-ERT2^* x *Ccr4^fl/fl^* mice exhibited a significant increase in eosinophils, Th2 cells, and mucus metaplasia (Figure 3G-J). We did not detect any difference in neutrophil numbers (Figure S4C) or in the frequency or numbers of Th1 cells, Th17 cells, or ILC2s within the lungs (Figures S4D-F). Of note, we also did not observe a difference in the Th2 cell or cDC2 populations in the mediastinal lymph node in our control and experimental groups (Figures S4G-L). Lastly, we did not detect any difference in serum IgE levels in *Foxp3^EGFP-Cre-ERT2^* x *Ccr4^fl/fl^* mice compared to controls (Figure S4M).

Given that allergic asthma in humans is a chronic inflammatory disease, we also characterized the role of CCR4 expression in Tregs during chronic allergic airway disease. We treated *Foxp3^EGFP-Cre-ERT2^* control mice and *Foxp3^EGFP-Cre-ERT2^* x *Ccr4^fl/fl^* mice with 10 μg of intranasal HDM 3 times per week for 6 weeks. We administered tamoxifen daily for five consecutive days between weeks 1-2, 3-4, and 5-6 of HDM exposure (Figure 3K). Compared to controls, *Foxp3^EGFP-Cre-ERT2^* x *Ccr4^fl/fl^* mice displayed selective deletion of CCR4 in Tregs (Figures 3L-M).

Similar to the acute model, deletion of CCR4 in Tregs did not reduce the percent or number of intraparenchymal Tregs but, rather, there was a dramatic increase in the total number of Tregs in the lungs on day 42 (Figure 3N, S4N). Despite the increased number of lung Tregs, *Foxp3^EGFP-Cre-ERT2^* x *Ccr4*^f*l/fl*^ mice exhibited a significant increase in the number of cDC2s in the lungs when compared to controls with no difference in cDC1s and only a modest increase in MCs (Figure 3O). Among lung cDC2s, there was a significant increase in the frequency and number of activated PD-L2^+^CCR7^+^ cDC2s (Figures S4O, 3P). Additionally, deletion of *Ccr4* in Tregs resulted in a significant increase in the number of Th2 cells, eosinophils, and mucus metaplasia in the lungs (Figures 3Q-R, S4P-Q). Taken together, these results indicate that CCR4 is not required for Treg trafficking into the lung parenchyma, but plays a critical role in suppressing the expansion of activated cDC2s, Th2 cells, and the severity of allergic inflammation.

### CCR4 promotes the efficiency of Treg trafficking to suppress the expansion of CCR7+ cDC2-CD4^+^ T cell clusters during allergic inflammation

Given our findings that CCR4 is not absolutely required for Treg trafficking into the lungs but serves a critical role in Treg suppression of allergic inflammation, we hypothesized that CCR4 is required for co-localizing Tregs with activated cDC2s. To define the role of CCR4 in regulating the positioning of Foxp3^+^ Tregs and activated cDC2s, we performed confocal imaging. We treated *FoxP3^EGFP-Cre-ERT2^* mice and *FoxP3^EGFP-Cre-ERT2^* x *Ccr4^fl/fl^* mice with HDM (days 0, 7-11) and tamoxifen (days 4-8). On day 14, we performed lung fixation and generated 250 µm thick sections, followed by staining for FSCN1, CD4, and Foxp3. FSCN1 is a well-characterized marker of CCR7^+^ cDCs, which we confirmed in our scRNA-seq dataset of lung DCs (Figure S5A).^57^ We performed tissue clearing and acquired 3D z-stack images. Compared to controls, lungs from *FoxP3^EGFP-Cre-ERT2^*x *Ccr4^fl/fl^* mice exhibited dramatically larger clusters of CD4^+^ T cells (Figure 4A). Magnification of the clusters revealed Foxp3^+^ cells within clusters containing CD4^+^ and FSCN1^+^ cells in both groups (Figure 4B). We performed a 3D spatial analysis to measure the distance of Foxp3^+^ Tregs and conventional CD4^+^ T cells from FSCN1^+^ surfaces within a 30 µm radius. Unexpectedly, we found that Tregs lacking CCR4 were positioned in similar proximity to FSCN1^+^ DCs as Tregs in the control group (Figure 4C). However, conventional CD4^+^ T cells were closer to FSCN1^+^ DCs in *Foxp3^-Cre-ERT2^* x *Ccr4^fl/fl^* mice than controls, consistent with greater DC-T cell clustering (Figure 4D). Our results suggest that CCR4 is not absolutely required for transmigration or co-localization with activated cDC2s with other chemokine receptors presumably compensating due to the greater inflammatory response.

**Figure 4.**
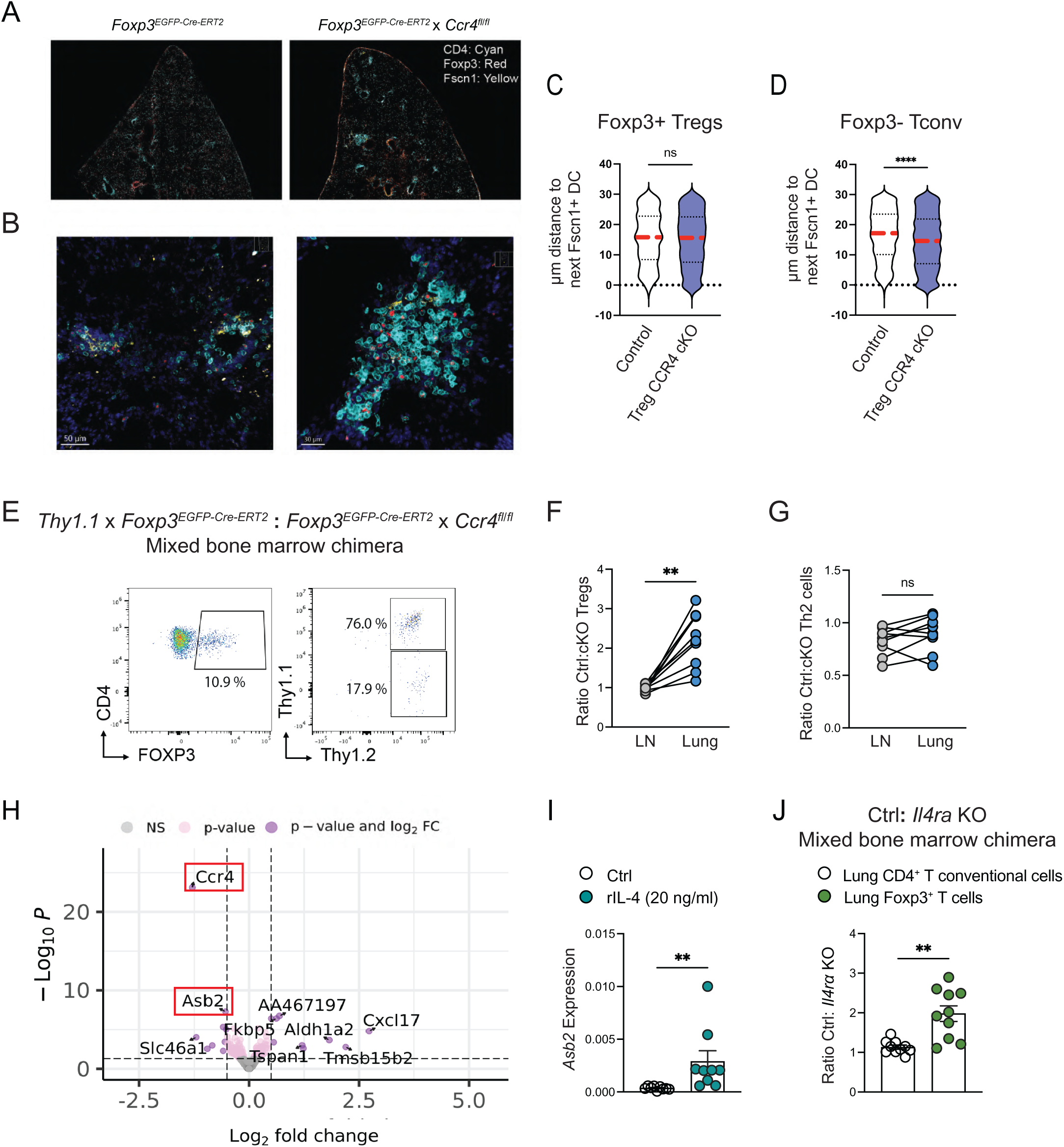
CCR4 promotes the efficiency of Treg trafficking to suppress the expansion of CCR7+ cDC2-CD4+ T cell clusters during allergic inflammation. *Foxp3^EGFP-Cre-ERT2^* control mice and *Foxp3^EGFP-Cre-ERT2^* x *Ccr4^fl/fl^* mice were sensitized with 10 µg i.n. HDM followed by daily challenges of 10 µg i.n. HDM on days 7–11, 1 mg daily doses of tamoxifen via oral gavage on days 4-8, then injected with anti-CD45 antibody i.v. 3 minutes prior to tissue harvest on day 14. **A.** Representative imaging of lungs from indicated groups with staining for CD4 (Cyan), Foxp3 (Red), and FSCN1 (Yellow). **B.** Higher magnification of images shown in (A). **C-D.** Quantification of nearest distance (µm) between FSCN1^+^ DCs and Foxp3^+^ Tregs (C) and FSCN1^+^ DCs and CD4^+^ T cells (D). **E-G**. Mixed bone marrow chimeras were generated with *Foxp3^EGFP-Cre-ERT2^*Thy1.1^+^Thy1.2^+^ control and *Foxp3^EGFP-Cre-ERT2^* x *Ccr4^fl/fl^* (Thy1.2) bone marrow cells. Ctrl:*Ccr4* cKO mixed bone marrow chimeras were treated with HDM (days 0, 7-11) and tamoxifen via oral gavage (days 4-8) then injected with anti-CD45 antibody i.v. 3 minutes prior to tissue harvest on day 14. **E.** Representative flow cytometry of lung CD4^+^ T cells showing Foxp3^+^ and Thy1.1^+^Thy1.2^+^ (Ctrl) and Thy1.2^+^ (*Ccr4* cKO) Tregs. **F.** Ratio of Ctrl:*Ccr4* cKO Tregs in the lymph nodes and lungs. **G.** Ratio of Ctrl:*Ccr4* cKO Th2 cells in the lymph nodes and lungs. **H.** Volcano plot showing differential gene expression analysis of bulk transcriptomes between sorted lung Tregs from *Foxp3^EGFP-Cre-ERT2^* control mice and *Foxp3^EGFP-Cre-ERT2^* x *Ccr4^fl/fl^* mice treated with HDM and tamoxifen. **I**. *Asb2* relative RNA expression in iTregs cultured with or without rIL-4 for 2 days. **J**. Mixed bone marrow chimeras were generated with C57BL/6 control (Ctrl) and *Il4r*α-deficient bone marrow cells. Ctrl:*Il4ra*-deficient mixed bone marrow chimeras were treated with HDM (days 0, 7-11) then injected with anti-CD45 antibody i.v. 3 minutes prior to tissue harvest on day 14. Ctrl:*Il4ra KO* ratio of lung T cell subsets. Images are representative of one of two independent experiments (A-B), data were pooled form two independent experiments (C-D), or data are representative of one experiment with *n*L=L10 mice per group from two independent experiments (E-G), or data were pooled form two independent experiments with *n* = 6 mice per group (H), or data were pooled from nine independent experiments (I), or data were pooled from two independent experiments with *n* = 10 mice (J). For statistical analysis, a two-tailed *t* test was performed for parametric data, and a two-tailed Mann-Whitney *U* test was performed for nonparametric data. **, p<0.001; ****, p<0.0001; ns, not significant.

These results suggested that deletion of *Ccr4* in Tregs either reduced the efficiency of Treg trafficking during the course of the Th2 cell response or impaired the ability of Tregs to locally suppress within cDC2-Th2 cell clusters or both.

To test the role of CCR4 in Treg trafficking efficiency, we generated mixed bone marrow chimeras with a 1:1 mix of *Foxp3^EGFP-Cre-ERT2^* (Thy1.1/1.2) and *Foxp3^EGFP-Cre-ERT2^* x *Ccr4^fl/fl^* (Thy1.2) bone marrow cells. We treated mixed bone marrow chimeras with HDM (days 0, 7-11) and tamoxifen (days 4-8). On day 14, we harvested lungs and mediastinal lymph nodes (mLNs) and found that control Tregs exhibited markedly greater efficiency in trafficking into the lung parenchyma compared to Tregs that deleted *Ccr4* (Figure 4E-F). Notably, there was no difference in Treg accumulation within the mLNs (Figure 4F). As a control, we compared the ratio of Th2 cells from both donor populations, which demonstrated similar accumulation within the mLN and lungs (Figure 4G). Taken together, our results indicate that Tregs require CCR4 to efficiently traffic into the lungs and suppress the expansion of cDC2-Th2 cells clusters during allergic inflammation.

### IL-4 receptor signaling promotes Treg trafficking in a cell-intrinsic manner

While our previous experiments show that Tregs require CCR4 to efficiently traffic into the lungs during allergic inflammation, we next investigated whether deleting *Ccr4* also impaired specific Treg suppressive mechanisms. To elucidate the role of CCR4 in regulating Treg function, we examined CCR4-dependent regulation of the Treg transcriptome. We treated *Foxp3^EGFP-Cre-ERT2^* mice and *Foxp3^EGFP-Cre-ERT2^* x *Ccr4^fl/fl^* mice with HDM (days 0, 7-11) and tamoxifen (days 4-8). On day 14, we sorted CD4^+^GFP^+^ Tregs from the lungs, isolated RNA, and performed bulk RNA-seq analysis. After quality control, we classified genes as differentially expressed based on a two-fold cutoff and false discovery rate <0.05. Differential expression analysis confirmed that *Ccr4* expression was significantly downregulated in Tregs from *Foxp3^EGFP-Cre-ERT2^* x *Ccr4^fl/fl^* mice compared to Tregs from *Foxp3^EGFP-Cre-ERT2^*controls (Figure 4H). Deletion of CCR4 in Tregs did not lead to reduced mRNA expression of canonical inhibitory molecules or suppressive cytokines such as IL-10 or TGF-beta, nor did we find any differences in the expression of genes regulated by TCR signaling. We also did not find any difference in the expression of STAT5-regulated genes to suggest differences in IL-2 sequestration. Unexpectedly, Tregs from *Foxp3^EGFP-Cre-ERT2^* x *Ccr4^fl/fl^* mice expressed significantly lower levels of the E3 ubiquitin ligase gene *Asb2* than control Tregs (Figure 4H). Asb2 is known to specifically target the actin-binding proteins filamin (FLN) A and FLNB for ubiquitylation and proteasomal degradation.^38^ FLNA and FLNB couple the actin cytoskeleton to integrins in their inactivate state, limiting integrin-dependent cell migration.^38^ Previous studies have shown that Th2 cells uniquely upregulate *Asb2* expression compared to other T cell subsets, which occurs in an GATA3-dependent manner.^58,59^ Loss of *Asb2* in Th2 cells leads to increased FLN expression, decreased integrin-dependent motility, and reduced allergic airway inflammation.^59^ Given that IL-4 promotes *Asb2* expression in Th2 cells, we next questioned whether CCR4 promotes efficient Treg trafficking towards niches high in IL-4, which directly induce Tregs to express *Asb2*.^59^ We cultured naive CD4^+^ T cells under Treg-polarizing conditions in vitro and after confirming successful Foxp3 expression, we stimulated cells with rIL-4. After 2 days, we detected a significant increase in *Asb2* expression in Tregs cultured in the presence of IL-4 compared to control conditions (Figure 4I). These findings led us to hypothesize that IL-4 directly enhances Treg trafficking efficiency during allergic inflammation in vivo. To test this hypothesis, we generated mixed bone marrow chimeras with a 4:1 mix of control and *Il4r*α KO bone marrow cells and treated mice with HDM (days 0, 7-11). At peak allergic inflammation, we assessed the ratio of control:*Il4r*α KO Tregs in the lungs. We found a significant increase in the ratio of control:*Il4r*α KO Tregs in the lungs but not in the ratio of conventional (Foxp3^-^ST2^-^) CD4^+^ T cells (normalized to spleen) (Figure 4J). Of note, control and *Il4r*α KO Tregs within the lungs exhibited no difference in Ki67 positivity, demonstrating similar proliferative potential (Figure S5B). These findings suggest that CCR4 and IL-4R signaling collaborate to promote Treg trafficking efficiency during allergic inflammation in a cell-intrinsic manner.

### Tregs require CCR4 to restrain co-stimulatory molecule expression in tissue cDC2s

While our bulk RNA-seq dataset of control and CCR4-deficient Tregs demonstrated that CCR4 promotes *Asb2* expression, the other differentially expressed genes exhibited low level expression, making their biological importance seem unlikely. In turn, we next investigated additional mechanisms whereby CCR4 may specifically control tissue cDC2 biology. Foxp3^+^ Tregs are known to suppress APC function via CTLA-4-dependent blockade or trans-endocytosis/trogocytosis of the co-stimulatory molecules CD80 and CD86.^60–63^ Furthermore, studies have shown that blocking CD80 and CD86 after Th2 cell priming reduces allergic inflammation, suggesting that co-stimulatory molecules promote effector Th2 cell responses.^64–66^ To test the role of CCR4 expression in Tregs in controlling CD80 and CD86 expression in lung APCs, we treated *Foxp3^EGFP-Cre-ERT2^* mice and *Foxp3^EGFP-Cre-ERT2^* x *Ccr4^fl/fl^* mice with HDM (days 0, 7-11) and tamoxifen (days 4-8). On day 14, we assessed the surface expression of CD80 and CD86 in lung cDC2s and MCs. Notably, compared to controls, lung cDC2s from *Foxp3^EGFP-Cre-ERT2^* x *Ccr4^fl/fl^* mice exhibited significantly higher expression of CD80 and CD86, whereas we found no difference in co-stimulatory molecule expression in MCs between the experimental groups (Figure 5A-D). We also assessed the ability of lungs Tregs to acquire CD80 and CD86 from APCs, finding that Tregs from *Foxp3^EGFP-Cre-ERT2^*x *Ccr4^fl/fl^* mice had lower CD80 and similar CD86 expression compared to control mice (Figure 5E-F). Given that cDC2s from *Foxp3^EGFP-Cre-ERT2^* x *Ccr4^fl/fl^* mice exhibit significantly more surface CD86 (Figure 5A), the similar expression of CD86 in Tregs between the two groups indicates CCR4 is required for efficient trans-endocytosis/trogocytosis (Figure 5E-F). In sum, our findings suggest that Tregs require CCR4 to specifically control co-stimulatory molecule expression in tissue cDC2s, but not MCs, during allergic inflammation.

**Figure 5.**
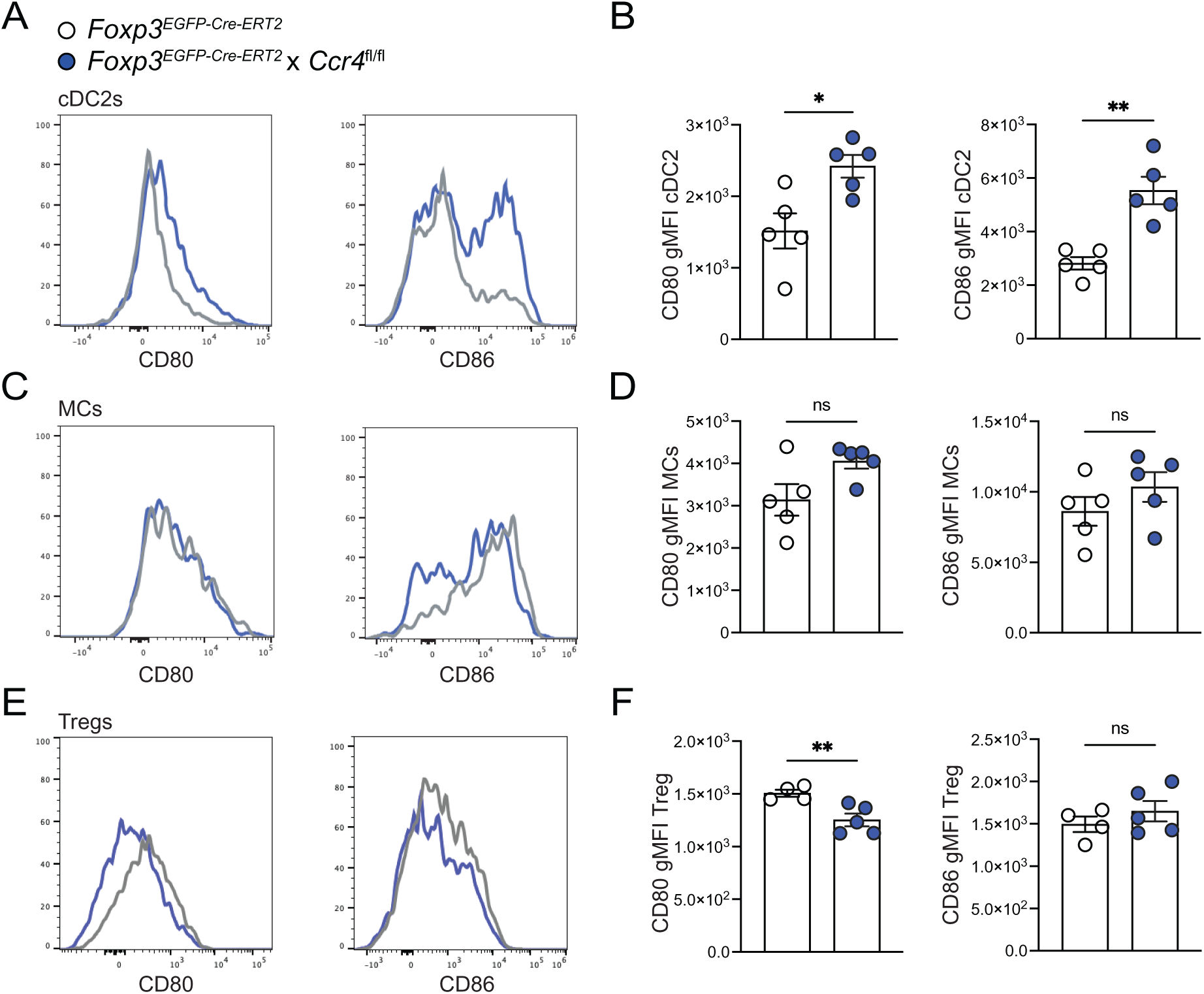
Tregs require CCR4 to specifically control cDC2 co-stimulatory molecule expression. *Foxp3^EGFP-Cre-ERT2^* control mice and *Foxp3^EGFP-Cre-ERT2^* x *Ccr4^fl/fl^* mice were sensitized with 10 µg i.n. HDM followed by daily challenges of 10 µg i.n. HDM on days 7–11, 1 mg daily doses of tamoxifen via oral gavage on days 4-8, then injected with anti-CD45 antibody i.v. 3 minutes prior to tissue harvest on day 14. **A.** Representative histograms of surface CD80 and CD86 expression on lung cDC2s from indicated groups. **B.** gMFI for surface CD80 and CD86 on lung cDC2s. **C.** Representative histograms of surface CD80 and CD86 expression on lung MCs. **D.** gMFI for CD80 and CD86 on lung MCs. **E.** Representative histograms of surface CD80 and CD86 expression on lung Foxp3^+^ Tregs. **F.** gMFI for CD80 and CD86 on lung Foxp3^+^ Tregs. Representative data shows individual mice with mean ± SEM from one of three independent experiments with *n*L=L4-5 mice per group. For statistical analysis, a two-tailed *t* test was performed for parametric data. *, p<0.05, **, p<0.01, ns, not significant.

### CCR4 ligand expression in activated cDCs in the human airway mucosa correlates with Treg enrichment

Our experimental work in mice suggests that expression of the CCR4 ligands in activated, tissue cDC2s promotes Treg trafficking efficiency. To test for the presence of an activated cDC2-Treg circuit in human airways during allergic inflammation, we leveraged a published, scRNA-seq dataset of airway mucosal cells in humans with allergy.^7^ Individuals with allergic asthma versus allergy without asthma were characterized as previously described.^7^ Specifically, recruited subjects had a clinical history of allergic conjunctivitis and/or rhinitis to HDM or cat, which was confirmed by skin prick testing with quantitative skin prick testing performed to standardize allergen dose. Subjects underwent bronchoscopy with baseline endobronchial brush samples collected at baseline and 24 hours after administration of diluent or allergen. scRNA-seq was performed on all live cells isolated from the brush. First, we re-analyzed the data to determine the expression of *CCL17* and *CCL22* across the cell lineages, finding that both CCR4 ligands are expressed in mononuclear phagocytes (MNPs), but not other cell types (Figure 6A). We next assessed the expression of *CCL17* and *CCL22* within MNP subsets, finding that *CCL17* is most highly expressed in DC2 (*CD1C*) and migDC (*CCR7*) clusters with *CCL22* exhibiting restricted expression in the migDC (*CCR7*) cluster (Figure 6B). The observed expression pattern for the CCR4 ligands in the human airways is remarkably similar to our murine dataset, suggesting that migDC (*CCR7*) cells promote the most chemotaxis for CCR4-expressing T cells (Figures 1D; S1C-D). We were not able to definitively determine the extent to which cells within the migDC (*CCR7*) cluster are derived from DC1 or DC2 cells. However, given that there is high expression of *CCL17* in the DC2 (*CD1C*) cluster and minimal expression of *CCL17* in the DC1 (*CLEC9A*) cluster suggests that during allergic inflammation the CCR4 ligands are preferentially expressed in tissue DC2 cells rather than DC1 cells (Figure 6B).

**Figure 6.**
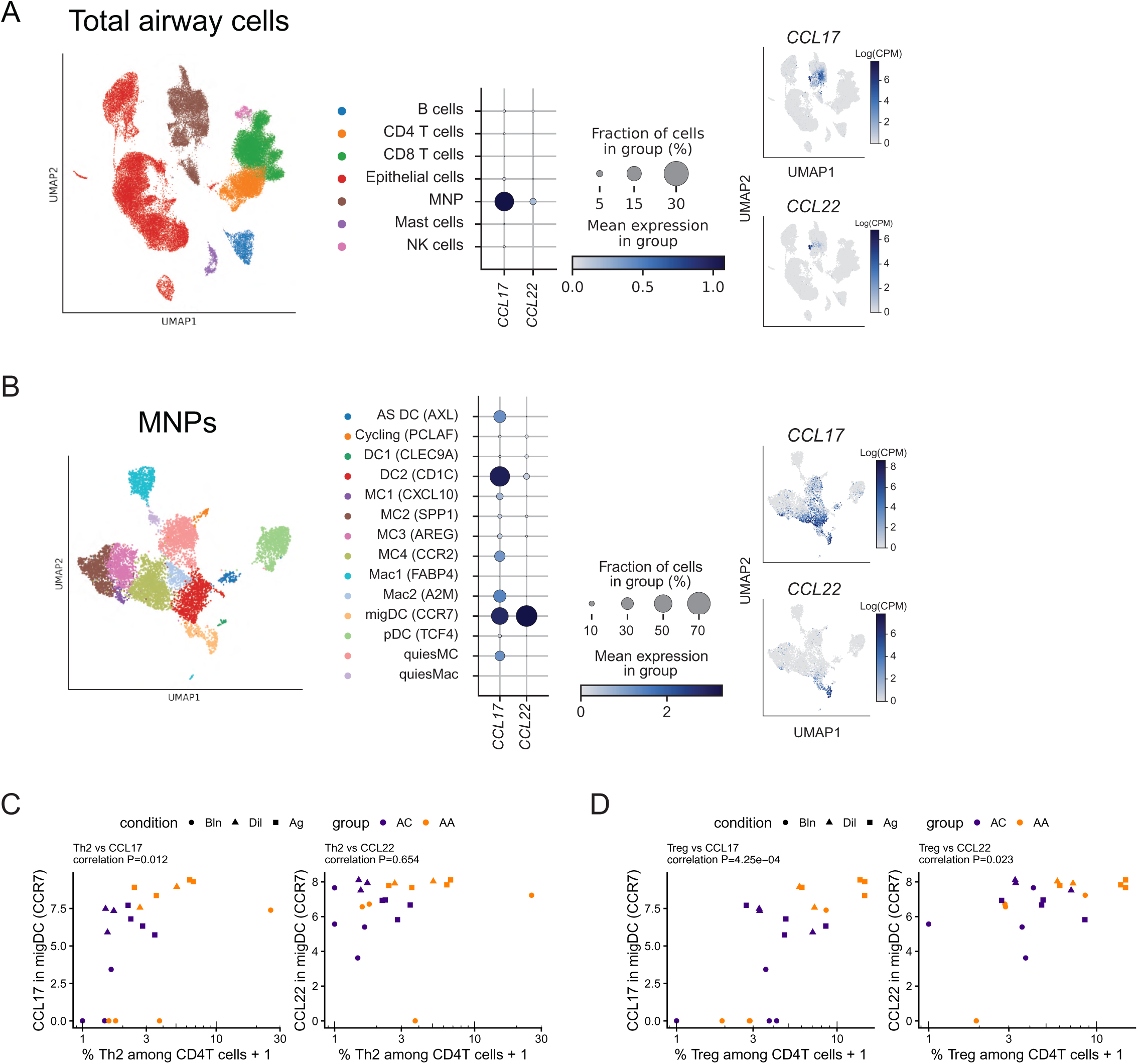
CCR4 ligand expression in activated cDCs in the human airway mucosal correlates with Treg enrichment. **A.** (left) UMAP embedding of single cells color-coded by predicted cell lineage with indicated annotation. (middle) Dot plot showing the percentage and expression level of *CCL17* and *CCL22* in each cell lineage with size of the dot representing the percentage of cells within each lineage expressing the chemokines and the color intensity indicating the scaled expression level. (right) Feature plot of *CCL17* and *CCL22* expression using pseudo-coloring to indicate gene expression. **B.** (left) UMAP embedding of MNPs color-coded predicted cell subsets with indicated annotation. (middle) Dot plot showing the percentage and expression level of *CCL17* and *CCL22* in each cluster with size of the dot representing the percentage of cells within each cluster expressing the chemokines and the color intensity indicating the scaled expression level. (right) Feature plot of *CCL17* and *CCL22* expression using pseudo-coloring to indicate gene expression. **C-D.** Correlation of *CCL17* and *CCL22* expression in migDC (*CCR7*) cluster (y-axis) with enrichment of Th2 cells (C) and Tregs (D) among CD4^+^ T cells (x-axis). P-values are from Pearson’s correlation test.

Lastly, we compared the expression of *CCL17* and *CCL22* in the migDC (*CCR7*) cluster with enrichment of Th2 cells and *FOXP3*-expressing Tregs in the airways. We found a correlation between *CCL17* expression, but not *CCL22* expression, in the migDC (*CCR7*) cluster with Th2 cell enrichment (Figure 6C). Notably, we found that both *CCL17* and *CCL22* expression in the migDC (*CCR7*) cluster correlated with the enrichment of Tregs in the airways (Figure 6D).

Overall, these findings are supportive of our proposed model in which activated cDC2s within the barrier tissue produce the CCR4 ligands to promote Treg chemotaxis to control Th2 cell-mediated inflammation.

## Discussion

Pathogenic effector Th2 cells play a critical role in driving allergic inflammation.^4–7^ In humans with allergic diseases, pathogenic effector Th2 cells are numerically rare within tissues, suggesting mechanisms that limit Th2 cell expansion at barrier sites.^7–13^ Along with pathogenic Th2 cells, Tregs and activated cDCs expressing the CCR4 ligands are enriched at barrier sites in chronic allergic diseases.^7–10^ However, the logic of the intercellular crosstalk regulating allergic immunity at barrier surfaces has remained unclear. Using a murine model of allergic lung inflammation, we demonstrate a feedforward circuit between effector Th2 cells and cDC2s in the lungs with the type 2 cytokines directly promoting cDC2 activation and the expression of the CCR4 ligands. Inducible deletion of *Ccr4* in Tregs led to a specific expansion of Th2 cells and activated cDC2s as well as increased allergic inflammation, suggesting that Tregs control the effector Th2 cell response by sensing tissue cDC2 activation via the CCR4 ligands.

Unexpectedly, we demonstrate that CCR4 is not absolutely required for Treg transmigration into the lungs or positioning within cDC2-CD4^+^ T cell clusters with other chemokine receptors presumably compensating. However, deletion of *Ccr4* in Tregs led to reduced trafficking efficiency into the lungs during allergic inflammation. CCR4 deficiency in Tregs did not impair canonical immunosuppressive cytokine expression, but did lead to reduced expression of the E3 ubiquitin ligase Asb2, which induces proteasomal degradation of filamin proteins to promote integrin-dependent cell migration.^38^ We show that IL-4 is sufficient to induce Asb2 expression in Tregs, suggesting that IL-4 instructs a pro-migration Treg state. We further demonstrate that Tregs lacking IL-4Rα exhibit diminished trafficking into the lungs during allergic immunity, suggesting CCR4 and IL-4R signaling collaborate to promote Treg trafficking to the sites of allergic immunity. Overall, we demonstrate that tissue cDC2s serve as a critical hub for effector Th2 cell and Treg function during allergic immunity with Treg suppression being tightly controlled by the magnitude of the effector Th2 cell response.

Our experimental work expands our insights into the role of cDC2s in allergic immunity. cDC2s are well established as critical initiators of Th2 cell priming in lymph nodes.^14–17^ Activated cDC2s traffic to the draining lymph node, uniquely generating cDC2-CD4^+^ T cell clusters that drive IL-2-and IL-4-dependent Th2 cell priming at the T-B border of the lymph node paracortex.^67,68^ Our results suggest that cDC2 regulation of Th2 cell responses extends beyond this initial priming event with tissue cDC2s controlling both effector Th2 cell and Treg function. Such an immunomodulatory role for cDCs has been described in the context of tumors. In tumors from humans and mice, CCR7^+^ cDCs are enriched in the tumor microenvironment with murine models suggesting a role in both promoting and restraining anti-tumor immunity.^43,69–73^ CCR7^+^ cDCs in the tumor microenvironment are derived from both cDC1s and cDC2s, produce IL-12, and recruit Tregs via CCL22.^43,72^ Our findings suggest that such “regulatory” cDC properties are not unique to the tumor microenvironment or IL-12-producing cDCs and that cDC2s can exhibit a phenotypic and functionally analogous regulatory role in non-lymphoid tissues during a type 2 immune response. While our study suggests that cDC2s are a critical tissue APC during allergic inflammation, the role of other tissue APCs in controlling the effector Th2 cell response remains unclear. Other studies have suggested that peribronchial, interstitial macrophages directly acquire inhaled antigen and can either directly present antigen epitopes to Th2 cells or pass antigen to cDC2s.^74^ ILC2s have also been shown to present allergen epitopes via MHCII expression to directly stimulate Th2 cells.^75^ Furthermore, additional APC subsets have been suggested to promote Th2 cell function.^76^ An integrated understanding of the role of various tissue APCs in regulating effector Th2 cells and Tregs during type 2 immunity remains an important area of future investigation.

Our study also raises important questions regarding how chemokines control allergic immunity in barrier tissues. Specifically, what are the rules dictating CCR7^+^ cDCs retention within the tissue versus migration via the afferent lymphatics to the draining lymph node?^43,69–73^ CCR7 binds to CCL19 and CCL21, which have different expression patterns and exhibit distinct biology. CCL19 is produced from adventitial and alveolar fibroblasts as well as smooth muscle cells and pericytes within the lungs.^77^ CCL21 is also produced by fibroblasts, but predominantly expressed by lymphatic endothelial cells and promotes cDC egress from barrier tissues to the draining lymph node.^78,79^ CCL21 possesses a highly charged C-terminal domain that binds glycosaminoglycans, generating a haptotactic gradient, whereas CCL19 lacks a charged C-terminus, resulting in free diffusion and generation of a classic chemotactic gradient.^80^ In tumors, CCL19-CCR7 interactions promote CCR7^+^ cDC positioning in perivascular niches within the tumor microenvironment, leading to the proposition that the relative signaling of CCL19 versus CCL21 dictates whether CCR7^+^ cDCs remain within a non-lymphoid tissue versus migrate to the draining lymph node.^72^ However, the mechanisms that may dictate relative sensing of the two CCR7 ligands during type 1 and 2 immune responses remain unclear. Along with the need for additional study in the regulation of the CCR7 system, our data raises important questions into the biology of biased agonism of the CCR4 system. While CCL17 and CCL22 both bind CCR4, they are known to exhibit distinct effects on CCR4. Specifically, in vitro studies have shown that CCL22 promotes greater recruitment of beta-arrestin to CCR4 and receptor internalization than CCL17.^81^ Along with this biased agonism, we show that CCL17 exhibits broader expression than CCL22 in tissue cDC2s in both mice and humans during allergic inflammation. Defining how such biased agonism and distinct expression patterns of CCL17 and CCL22 effect Th2 cell and Treg biology will require the development of novel tools to rigorously interrogate the CCR4 ligands individually in an inducible manner in vivo.

Beyond cDC2 and chemokine biology, our findings have important implications for our understanding of Treg biology during type 2 immunity. IL-4 has been shown to suppress Treg differentiation or function as well as promote an ex-Treg program that contributes to chronic allergic disease.^82–87^ These results have led to the view that IL-4 signaling inhibits Treg suppression. Other studies have suggested that IL-4 signaling in Tregs can promote their suppressive function, although the mechanistic basis has remained unclear.^88–90^ We demonstrate that type 2 cytokines enhance Treg trafficking efficiency by 1) promoting cDC2 expression of the CCR4 ligands and 2) directly stimulating Tregs. The mechanisms whereby IL-4R signaling promotes Treg trafficking versus suppresses or de-stabilizes the Treg program to generate ex-Tregs remains unclear and represents an important area for future study. The ability of the type 2 cytokines to promote Treg suppression of allergic immunity has important implications for novel therapeutic approaches for chronic allergic diseases. Notably, current therapeutic approaches for chronic allergic diseases aim to suppress type 2 cytokine production, but our work suggests these interventions are concomitantly attenuating Treg suppression and may limit approaches that aim to re-establish immune tolerance. In addition, there is growing interest in novel platforms to generate Treg-based cellular therapies to treat chronic inflammatory diseases.^91^ Our findings suggest that programing allergen-specific Tregs to be more sensitive to the CCR4 ligands and/or IL-4 signaling without inducing a pathogenic ex-Treg state would yield more effective suppression of chronic allergic inflammation.

In summary, we rigorously define the crosstalk between effector Th2 cells, cDC2s, and Tregs in a barrier tissue during allergic immunity. We propose a novel model whereby tissue cDC2s serve as a cellular platform controlling the magnitude of allergic inflammation by dictating the efficiency of Treg suppressive function with important implications for future investigation into type 2 immunity as well as the development of effective Treg-based therapies to durably suppress chronic allergic diseases.

### Limitations of the study

We acknowledge that our genetic approach to delete *Ccr4* in Foxp3^+^ Tregs results in loss of CCR4 expression in all Foxp3^+^ Tregs rather than specifically in allergen-specific Foxp3^+^ Tregs. Consequently, we are unable differentiate the role of CCR4 in allergen-specific versus self-specific Tregs. Nevertheless, for the first time, our approach allows for the inducible and selective deletion of *Ccr4* in Foxp3^+^ Tregs in vivo while maintaining CCR4 in other T cell subsets, including Th2 cells. In addition, we propose that CCR7^+^ cDC2s serve a critical role in the lungs during allergic immunity. While inducible depletion of activated cDC2s would allow for the assessment of cDC2 function in vivo, there is currently a lack of specific and inducible deletion models for CCR7^+^ cDC2s. In the future, novel experimental tools will be needed to gain selective and inducible genetic control of activated cDC2s in vivo.

### Experimental models and study participant details Mice

Mice were bred and maintained in a specific-pathogen-free (SPF) animal facility at Massachusetts General Hospital. C57BL/6J, CD45.1 CRL, MHCII-deficient (B6.129S2-*H2^dlAb1-Ea^*/J, stock no. 003584)^92^ and Thy1.1 (B6.PL-*Thy1^a^*/CyJ, stock no. 000406) mice were purchased from Charles River Laboratories (Wilmington, MA) or the Jackson Laboratory (Bar Harbor, ME). *Il4r ^-/-^* were provided by S. Bromley (Massachusetts General Hospital). *Ccr4* floxed mice were generated by the Rahimi lab with the assistance of Ingenious Targeting Laboratory with *loxP* sites introduced around exon 2 of the *Ccr4* locus via homologous recombination. *Ccr4* floxed mice were crossed to *Foxp3^EGFP-Cre-ERT2^* mice (*Foxp3^EGFP-Cre-ERT2^*; stock no. 016961)^56^ purchased from the Jackson Laboratory to generate *Foxp3^EGFP-Cre-ERT2^* x *Ccr4* floxed mice. C57BL/6J and CD45.1 CRL mice were crossed to generate CD45.1/CD45.2 mice. *Foxp3^EGFP-Cre-ERT2^* were crossed to Thy1.1 to generate *Foxp3^EGFP-Cre-ERT2^* x *Thy1.1* mice. Age and sex matched mice were used in experiments at 6-12 weeks of age. Both female and male mice were used for experiments and numbers were matched among the experimental conditions when possible. Animals were housed in a facility with a standard 12-hour light cycle and maintained at a standard temperature of 18–23L°C and humidity of 40–60% with accessible food and water. All animal experiments were approved by the Massachusetts General Hospital Subcommittee on Research Animal Care.

### Mouse treatments

To induce allergic airway inflammation, mice were anesthetized with intraperitoneal injection of ketamine/xylazine (Patterson Veterinary) and sensitized via i.n. administration with 10 µg HDM (Dermatophagoides pteronyssinus extracts; Greer Laboratories) in 40 µl sterile PBS on day 0. On day 7, mice were challenged with 10 µg HDM via i.n. route daily for 5 days. To block IL-4 and IL-13, the antibodies were injected intraperitoneally on days 9, 11 and 13 of HDM challenge at a dose of 0.25 mg per mouse for anti-IL-4 antibody, 0.1 mg per mouse for anti-IL-13 antibody, as well as 0.35 mg per mouse for isotype control (IgG1). To delete *Ccr4* in Foxp3^+^ Tregs, control and *Foxp3^EGFP-Cre-ERT2^* x *Ccr4^fl/fl^* were treated with tamoxifen (1 mg per mouse in 100 μl mixture of corn oil/ethanol) daily via oral gavage on days 4-8 of HDM treatment.

### Bone Marrow Chimera Generation

To generate bone marrow chimeras, 6-week-old CD45.1^+^ C57BL/6 recipient mice were irradiated with a split dose of 1000 rad (two doses of 500 rad, 24 hours apart). Donor bone marrow (BM) was extracted from femurs and tibias of mice. BM cells were incubated in red blood cell lysis buffer (Sigma-Aldrich), then the purified BM cells were resuspended in 1x PBS. 5 × 10^6^ BM cells in 100 μL of PBS were intravenously injected into CD45.1 C57BL/6 recipients within 18 h of final irradiation. For mixed chimeras, recipients received either a 1:4 ratio of control:*Il4rα^-/-^ BM or a 1:1 ratio of Foxp3^EGFP-Cre-ERT2^ X Thy1.1 and Foxp3^EGFP-Cre-ERT2^* x *Ccr4^fl/fl^* BM. Recipient mice were rested for 8 weeks before induction of the relevant HDM model. Lungs, mLN, and spleens were harvested for chimeric assessment.

### Tissue harvest and leukocyte preparation

Prior to tissue harvest, mice were anesthetized with ketamine/xylazine (Patterson Veterinary) and injected i.v. with 3 µg fluorophore-labeled anti-CD45 antibody (30-F11; BioLegend) through the retro-orbital sinus to label intravascular leukocytes. 3 min after anti-CD45 antibody injection, mice were euthanized and tissues were harvested. Lung lobes and mLNs were removed, minced with scissors, and digested at 37°C for 25 min in digestion buffer (0.52 U/ml Liberase TM (Roche) and 60 U/ml DNase I (Roche) in RPMI 1640 (Cellgro) with 5% FBS). Digested tissue was strained through a 70-µm filter followed by incubation with red blood cell lysis buffer (Sigma-Aldrich) to generate a single-cell suspension. Spleens were collected from the same mice as indicated and mechanically dissociated through a 70-μm filter and subjected to red blood cell lysis to generate a single-cell suspension.

### Generation of Bone Marrow-derived Dendritic Cells (BMDCs)

BM was harvested from 6-8-week-old mice, followed by incubation with red blood cell lysis buffer (Sigma-Aldrich). BM cells were cultured in cRPMI supplemented with 20 ng/mL of granulocyte-macrophage colony-stimulating factor (GM-CSF) (Peprotech) at a density of 1 x 10^6^ cells/mL and fed on day 3. On day 6, the non-adherent cells (granulocytes) were discarded, the loosely adherent cells collected with gentle pipetting, and the adherent cells collected using trypsin-EDTA (Sigma-Aldrich). Cells were re-plated in media containing GM-CSF (20 ng/ml) and left untreated or stimulated with IL-4 (Peprotech) at 20 ng/ml. Cells were incubated at 37°C with 10% CO_2_Lfor a further 4 days, adding media on day 8. At the end of the assay on day 10, cells were collected for flow cytometry or washed with PBS and lysed with Buffer RLT + 1% β-mercaptoethanol and RNA extracted using the Qiagen RNeasy Mini Kit as per the manufacturer’s instructions.

### In vitro generation of Tregs

Naïve CD4+ T cells were isolated from spleens of mice using a CD4 T cell isolation kit, according to the manufacturer’s instructions (STEMCELL Technologies). The isolated cells were cultured under Treg polarizing conditions (1ug/ml anti-CD3 coating, 1 ug/ml anti-CD28, 10 ug/ml anti-IFN γ, 5 ng/ml TGFβ and 100 U/ml IL-2. Naïve CD4 T cells were polarized under these conditions for 3 days, then cultured for an additional 2 days in the presence or absence of IL-4 (20ng/ml). Cells were cultured in cRPMI 1640 media (Cellgro) with 10% (vol/vol) heat inactivated fetal bovine serum (FBS, Sigma), 50 µM 2-mercaptoethanol, 2 mM Glutamax (Gibco), 5 mM Hepes, 1 mM sodium pyruvate, 0.1 mM nonessential amino acids, and 100 U/ml penicillin-streptomycin (all from Lonza).

### Flow cytometry and cell sorting

Staining was performed using the following fluorochrome-conjugated anti-mouse monoclonal antibodies purchased from Thermo Fisher, BD Pharmingen or BioLegend: CD45-APCeF780 (30-F11), SiglecF-BUV395 (E50-2440), Ly6G-AF700 (1A8), CD11c-BV605 (N418), MHCII-BUV737 (M5/114.15.2), SirpA-AF700 (P84), XCR1-BV650 (ZET), CD26-BV785 (H194-112), CD64-APC (X54-5/7.1), CCR7-biotinylated (4B12), PD-L2-BV421 (Ty-25), CD80-BV711 (16-10A1), CD86-BV510 (PO3), CD45.1-PerCpCy5.5 (A20), CD45.2-APC (104), Thy1.1-APC (O-X7), Thy1.2-BUV395 (532.1), CD3-BUV395 (UCHT1), CD4-BUV737 (RM-45), ST2-BV421-(U29-93), CCR4-biotinylated (2G12), CXCR3-BV650 (173), CCR6-BV605 (29-2L17), CD44-APC (IM7), ICOS-AF700 (C398.4A), PD-1-PerCP-Ef710 (J43), TIGIT-BV421 (1G9), GITR-PeCy7 (DTA-1), Ki67-BV785, GATA3-PE (L50-823), Foxp3-AF488 (FJK-16s) and Streptavidin-PE.

Dead cells were identified using Fixable Viability Dye eF780 or eF506 (eBioscience). For surface staining, single-cell suspensions were subjected to Fc-receptor blockade (clone 93; Fc Block, BioLegend) and were incubated for 30 min at 4 °C with the indicated mouse antibody. CCR4 and CCR7 staining were performed by incubating cells for 1 h at 37 °C with biotinylated antibodies. Cells were then washed and subjected to viability dye and surface staining. For transcription factor staining, cells were stained with viability dye and surface markers as documented above. Cells were then fixed and permeabilized with the eBioscience Foxp3/Transcription Factor Staining buffer set according to the manufacturer’s instructions (Thermo Fisher) and stained with indicated antibodies overnight at 4°C. Ab-labelled cells were analyzed on a Cytek Aurora (Cytek Biosciences) or sorted on a MA900 multi-application cell sorter (SONY, Cell Sorter Software v.3.3.2). Flow cytometric data were acquired using SpectroFlo v.3.2.1 (Cytek Biosciences) software, and.fcs files were analyzed using FlowJo v.10 (TreeStar).

### qPCR

Total RNA was extracted from tissues using the TRIzol/chloroform method. RNA was isolated from cells using the RNeasy Mini kit according to the manufacturer’s instructions (QIAGEN, 74106). cDNA was reverse transcribed using SuperScript III First Strand (Invitrogen) following manufacturer’s guidelines. Quantitative PCR (qPCR) reactions were performed on a LightCylcer 96 Instrument (Roche) using FastStart Essential DNA Green Master (Roche) and normalized to B2M using the following primers: *Ccl17*, 5’-CAG GGATGCCATCGTGTTTC-3’ (forward) and 5’-CACCAATCTGATGGCCTTCTT-3’ (reverse); *Ccl22*, 5’-TAC ATCCGTCACCCTCTGCC-3’ (forward) and 5’-CGGTTATCAAAACAACGCCAG-3’ (reverse); *Asb2*, 5’-CACTCTGGCTCTGCACCTTC-3’ (forward), 5’GGGCTCTGCAAGATTCTTCC3’ *B2m*, 5’-CCCGTTCTTCAGCATTTGGA-3’ (forward) and 5’-CCGAACATACTGAACTGC TACGTAA-3’ (reverse).

### Single cell RNA-seq

A single-cell suspension of lung cells was generated from pooled mice with 2 biological replicates of naive mice and 3 biological replicates of HDM-treated mice with 5 mice per replicate across 2 independent experiments. Lung intraparenchymal CD11c^+^MHCII^+^ cells were sorted into supplemented phenol free RPMI and 50% FBS. After washing the sorted cells, each population was loaded as a separate channel on the Chromium 10X Genomic v1.1 platform (10X Genomics). Libraries were prepared using the Chromium Single Cell 5’ Library Construction Kit. cDNA libraries were generated from the complementary DNA produced through Chromium chemistry. Sequencing of gene expression libraries was performed using the NextSeq 500/550 High Output kit v.2.5 (75 cycles; Illumina). Sequencing was performed on the NextSeq 550 (Illumina).

Raw sequencing data were processed with Cell Ranger 7.1.0 (10x Genomics). The pipeline converted Illumina basecall files to FASTQ format, and reads were aligned to the GRCm39 (mm39) mouse reference genome to generate a gene–cell count matrix. After Cell Ranger quality control, we retained 26,456 cells from 5 samples and merged them into a single Seurat object.^93,94^ Cells were filtered based on ≥500 detected genes, ≥1,000 UMIs, mitochondrial read fraction ≤10%, and a novelty score (log10[genes]/log10[UMIs]) ≥0.8 yielding a final dataset of 23,592 final cells. Following normalization and identification of variable features, we performed principal component analysis (PCA) on the merged object and selected the first 20 principal components. We then integrated the 5 samples using Seurat’s IntegrateLayers function with the CCAIntegration method (orig.reduction = “pca”, new.reduction = “integrated.cca”) to correct for sample-specific effects. The integrated reduction was used to construct a shared nearest-neighbor graph and to perform clustering with the Louvain algorithm at a resolution of 0.5, followed by computation of Uniform Manifold Approximation and Projection (UMAP) embeddings for visualization. Cluster identities (cell types) were assigned using a combination of manual inspection of canonical marker genes and differential expression analysis with FindMarkers().

### Bulk RNA-seq

A single-cell suspension of lung cells was generated from a total of 6 *Foxp3^EGFP-Cre-ERT2^*controls and 6 *Foxp3^EGFP-Cre-ERT2^* x *Ccr4^fl/fl^*mice across two independent experiments with tissue collected on day 14 of the HDM and tamoxifen treatment protocol.

Lung intraparenchymal Foxp3^+^ Tregs were sorted into Buffer RLT + 1% β-mercaptoethanol (∼25,000 cells/lung). Total RNA was extracted with a RNeasy Mini kit according to the manufacturer’s instructions (QIAGEN, 74106). Library construction and sequencing was carried out using the Novogene full Eukaryotic mRNA-seq pipeline. Combat-seq was used to correct for batch effect and DESeq2 was used to identify differentially expressed genes.

Raw FASTQ files were processed with nf-core/rnaseq (v3.14.0).^95^ Reads were aligned to the *Mus musculus* GRCm39 (mm39) genome with Salmon (v1.10.1) for differential expression.^96^ QC metrics (library size, mapping rate, etc.) were evaluated in R 4.3.1 using bcbioR rnaseq-reports (v0.4.5).^97^ Salmon gene-level counts were batch-corrected with ComBat-seq using sorting date as the batch variable and analyzed with DESeq2 v1.42.1.^98,99^ Genes with false discovery rate (FDR; Benjamini–Hochberg–adjusted P value) < 0.05 and |log_₂_ fold change| ≥ 1 were considered differentially expressed. Gene set enrichment analysis (GSEA) was performed with fgsea v1.28.0 using genes ranked by log_₂_ fold change and Gene Ontology (GO) gene sets obtained from MSigDB via msigdbr v7.5.1.^100–103^ Over-representation analysis (ORA) of significantly differentially expressed genes was carried out with clusterProfiler v4.10.1 using GO Biological Process, Cellular Component, and Molecular Function categories; GO terms with FDR < 0.05 were considered significantly enriched.^104^

### Confocal Microscopy

Lungs were perfused via the trachea with 2% PFA, fixed in 2% PFA for 2 h, and subsequently dehydrated in a graded sucrose gradient (10% and 20% for 2 hours each, and 30% overnight) at 4 °C. Tissues were washed and embedded in 1.5% low-melting-point agarose (Invitrogen) in PBS, and 250 µm thick consecutive sections were generated using a vibratome. Sections were transferred to a 24-well plate containing PBS with 0.1M glycine. Sections were permeabilized and blocked in PBS containing 10% BSA, Fc block (BioLegend), and 0.2% Triton X-100 overnight at 4°C. Primary antibody staining was performed in two rounds with anti-CD31 MEC13 (Biolegend), in blocking buffer for 48 hours at 4°C followed by Anti-Rat DyLight 755 (Invitrogen). Following a washing cycle in 0.2% Triton X-100/PBS, sections were incubated with secondary antibodies anti-CD4 BV480 (BD), anti-CD11c AF488 (Invitrogen), anti-FOXP3 efluor570 (Invitrogen) and anti-Fascin1 AF594 (Santa Cruz Biotechnology) for 48 hours. Following a washing cycle, tissues were stained with DAPI for 2 hours in PBS at room temperature and washed three times with PBS. To achieve tissue transparency for deep-tissue imaging, sections were incubated in RapidClear 1.47 for 48 hours under continuous agitation. Image analysis was performed using the IMARIS software v9.7 (Oxford Instruments). The IMARIS surface generation tool was used for cell rendering to visualize representative 3D objects for individual FSCN1^+^ DCs, and spots for Foxp3^+^ Treg and CD4^+^ T conventional cells. Distances between FSCN1^+^ DCs and T cell subsets were determined using the object-object shortest distance function in IMARIS.

### ELISA

Blood was collected from mice at time of harvest via cardiac puncture. Serum IgE was measured using a standard sandwich ELISA. Samples were assessed on a SpectraMax iD5 microplate reader (Molecular Devices).

### Histology

Lung samples were fixed in buffered 10% formalin solution. Paraffin-embedded sections were cut (5 mm) and stained with periodic acid–Schiff (PAS). Goblet cells were enumerated from PAS-stained sections using a numerical scoring system. 20–50 airways per mouse were evaluated, and the sum of goblet cell scoring from each mouse was divided by the number of airways and presented as a mucus score as previously described.^2^ All images are shown at 10X magnification.

## Statistical methods

Results are shown as meanL±LSEM. Symbols represent individual experimental replicates or mice or as described in figure legends. Unless otherwise noted, data were processed in Excel v.16.86 and statistical analyses were performed using Prism v.10 (GraphPad). When two or more normality tests indicated that the distribution of data was not normal, the appropriate nonparametric test was performed. When comparing one variable among three or more groups or two variables, analysis of variance (ANOVA) was always used. As indicated in the figure legends, two-sidedL*t*-tests, two-sided Mann–WhitneyL*U*-tests, one-way ANOVA with Tukey’s multiple comparisons tests, two-way ANOVA with Tukey’s multiple comparisons (comparing all groups) or two-way repeated measures ANOVA (with paired data) were used throughout the study. Paired analyses were used when samples were derived from the same subject. Data were significant if they met the followingL*p*Lvalue criteria: *, p<0.05; **, p<0.01; ***, p<0.001;****, p<0.0001; ns, not significant.

---

**Supplementary Figure 1.**
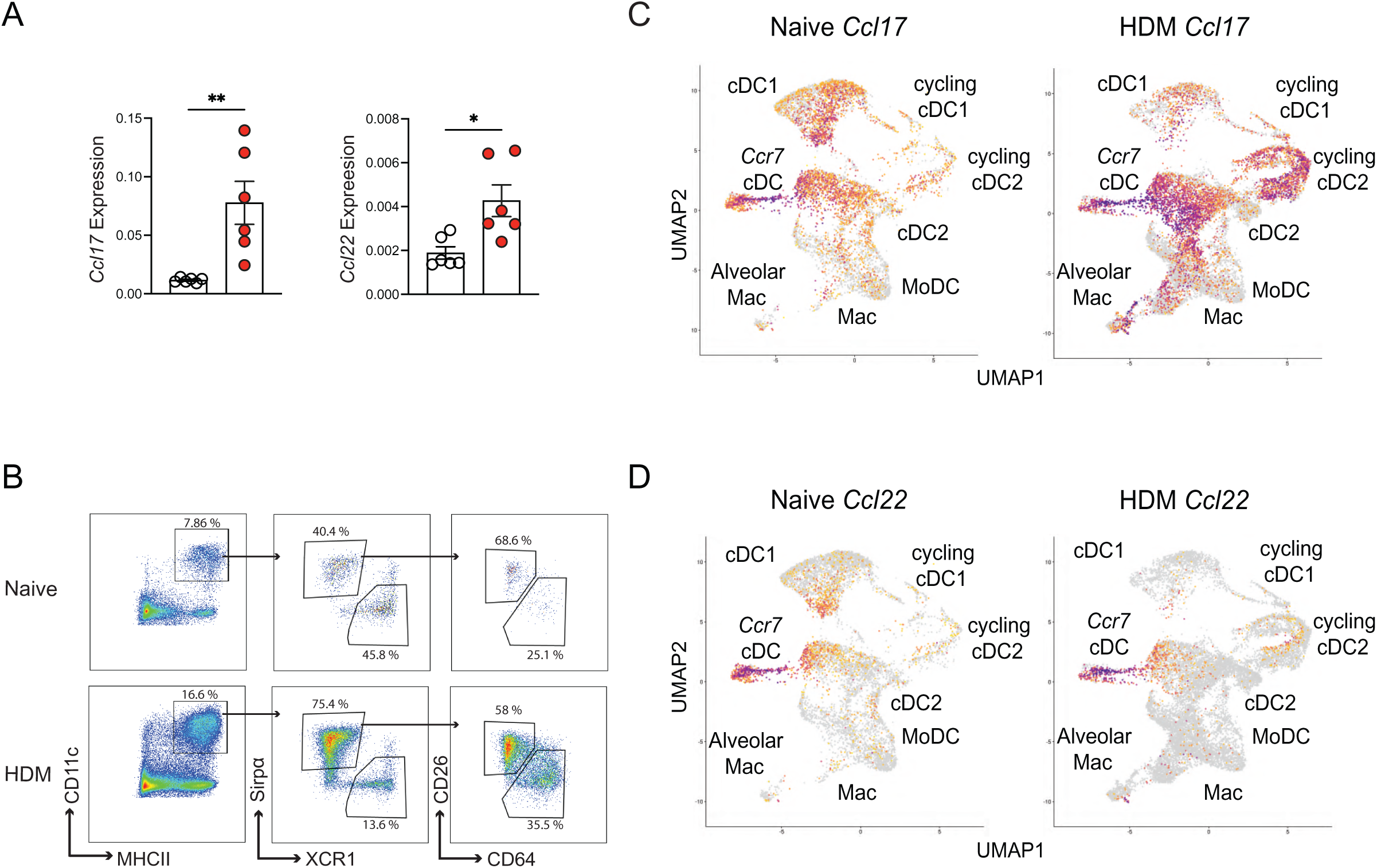
Characterization of the expression of the CCR4 ligands during allergic inflammation. C57BL/6 mice were sensitized (day 0) with 10 µg i.n. HDM followed by daily challenges of 10 µg i.n. HDM on days 7–11 then injected with anti-CD45 antibody i.v. 3 minutes prior to tissue harvest on day 14. **A.** Total lung *Ccl17* and *Ccl22* relative RNA levels assessed via qPCR. **B.** Representative flow cytometry and gating strategy for lung DC subsets. **C-D.** scRNA-seq UMAP of *Ccl17* and *Ccl22* expression in lung DC clusters from nailJve and HDM-treated mice. Representative data shows individual mice with mean ± SEM from one of three independent experiments with nL=L6 mice per group (A) or pooled from two independent experiments, *n* = 10 mice per group (C-D). For statistical analysis, a two-tailed *t* test was performed for parametric data. *, p<0.05; **, p<0.01.

**Supplementary Figure 2.**
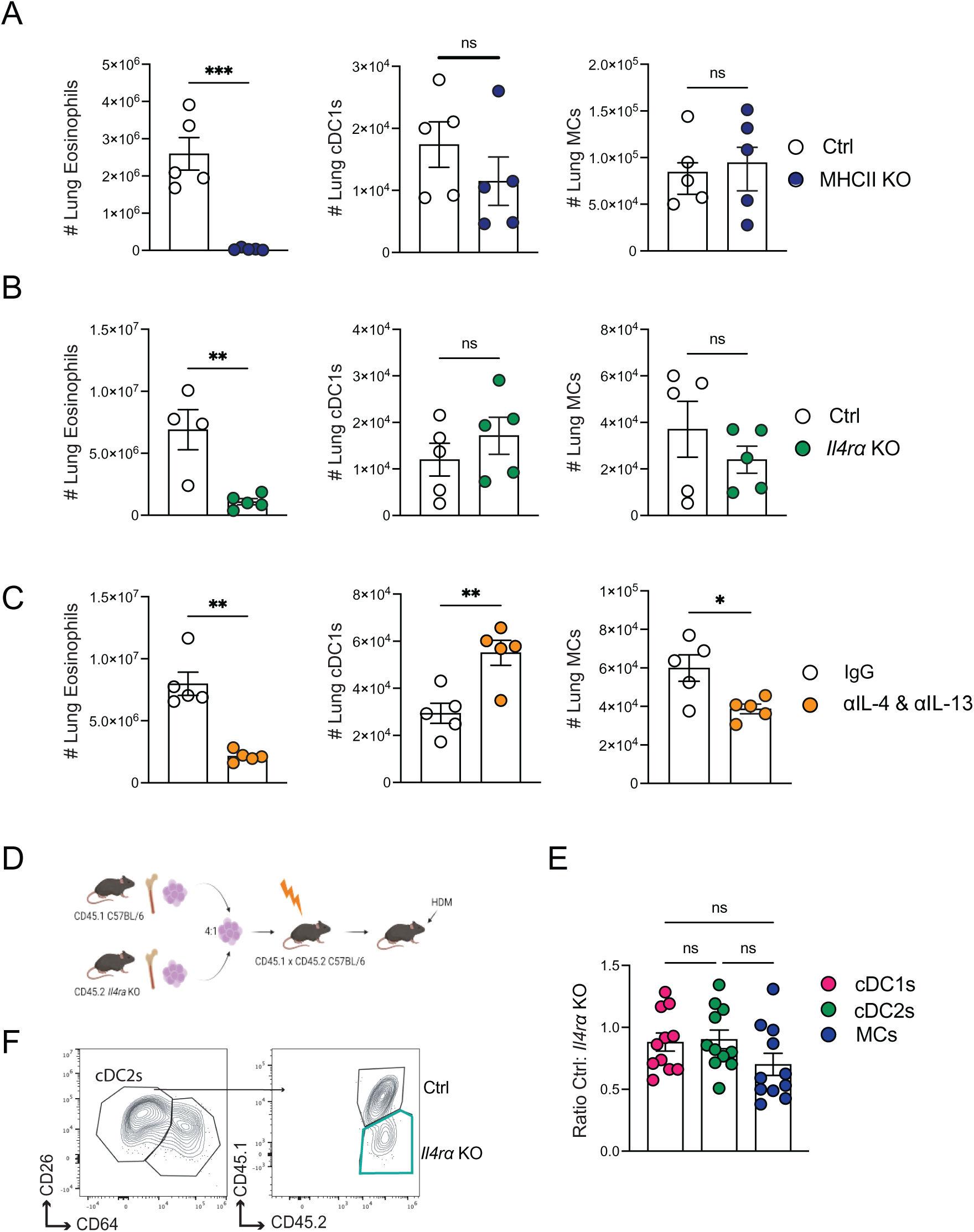
Defining the role of CD4^+^ T cells and type 2 cytokines in lung DC activation. Indicated mice were sensitized (day 0) with 10 µg i.n. HDM followed by daily challenges of 10 µg i.n. HDM on days 7–11 then injected with anti-CD45 antibody i.v. 3 minutes prior to tissue harvest on day 14. **A**. Number of total lung eosinophils, cDC1s, and MCs from HDM-treated Ctrl and MHCII KO mice. **B**. Number of total lung eosinophils, cDC1s, and MCs from HDM-treated Ctrl and *Il4ra* KO mice. **C**. C57BL/6 mice were treated as in (A-B) with administration of IgG or anti-IL-4/IL-13 antibodies via intraperitoneal (i.p) injection on days 7,11, 13. Number of total lung eosinophils, cDC1s, and MCs from indicated groups. **D**. Schematic of generation of Ctrl:*Il4ra KO* mixed bone marrow chimera mice. **E.** Ctrl:*Il4ra* KO ratio of indicated lung DC subsets. **F.** Representative flow cytometry of gating strategy for sorting Ctrl and *Il4ra* KO lung cDC2s from mixed bone marrow chimera mice. Data are representative of one experiment with nL=L5 mice per group from three independent experiments (A-C) or data were pooled from two independent experiments, *n* = 11 mice (E). For statistical analysis, a two-tailed *t* test was performed for parametric data. One-way ANOVA analysis with Holm-Sidak’s testing was performed for multiple comparisons. *, p<0.05; **, p<0.01, ***, p<0.001, ns, not significant.

**Supplementary Figure 3.**
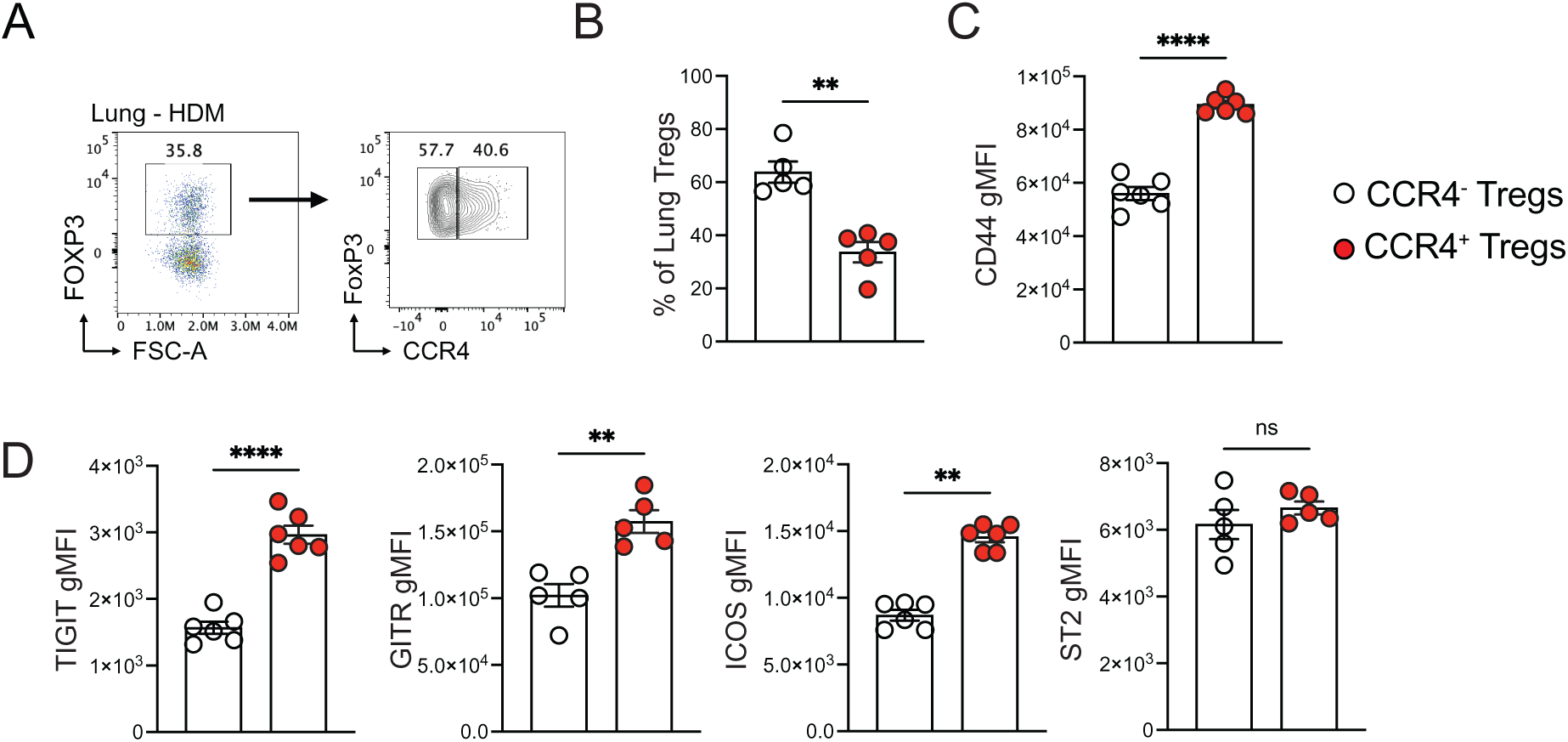
Characterizing the phenotype of CCR4^+^ Tregs during allergic inflammation. Mice were sensitized (day 0) with 10 µg i.n. HDM followed by daily challenges of 10 µg i.n. HDM on days 7–11 then injected with anti-CD45 antibody i.v. 3 minutes prior to tissue harvest on day 14. **A**. Representative flow cytometry of lung intraparenchymal Tregs showing CCR4 expression during allergic inflammation. **B**. Percent CCR4^-^ and CCR4^+^ Tregs during allergic lung inflammation. **C-D**. Expression of lung Treg markers CD44, TIGIT, GITR, ICOS, and ST2 in CCR4^-^ and CCR4^+^ Tregs during allergic inflammation. Data are representative of one experiment with *n*L=L5-6 mice per group from three independent experiments. For statistical analysis, a two-tailed *t* test was performed for parametric data and a two-tailed Mann-Whitney *U* test was performed for nonparametric data. **, p<0.001; ****, p<0.0001, ns, not significant.

**Supplementary Figure 4.**
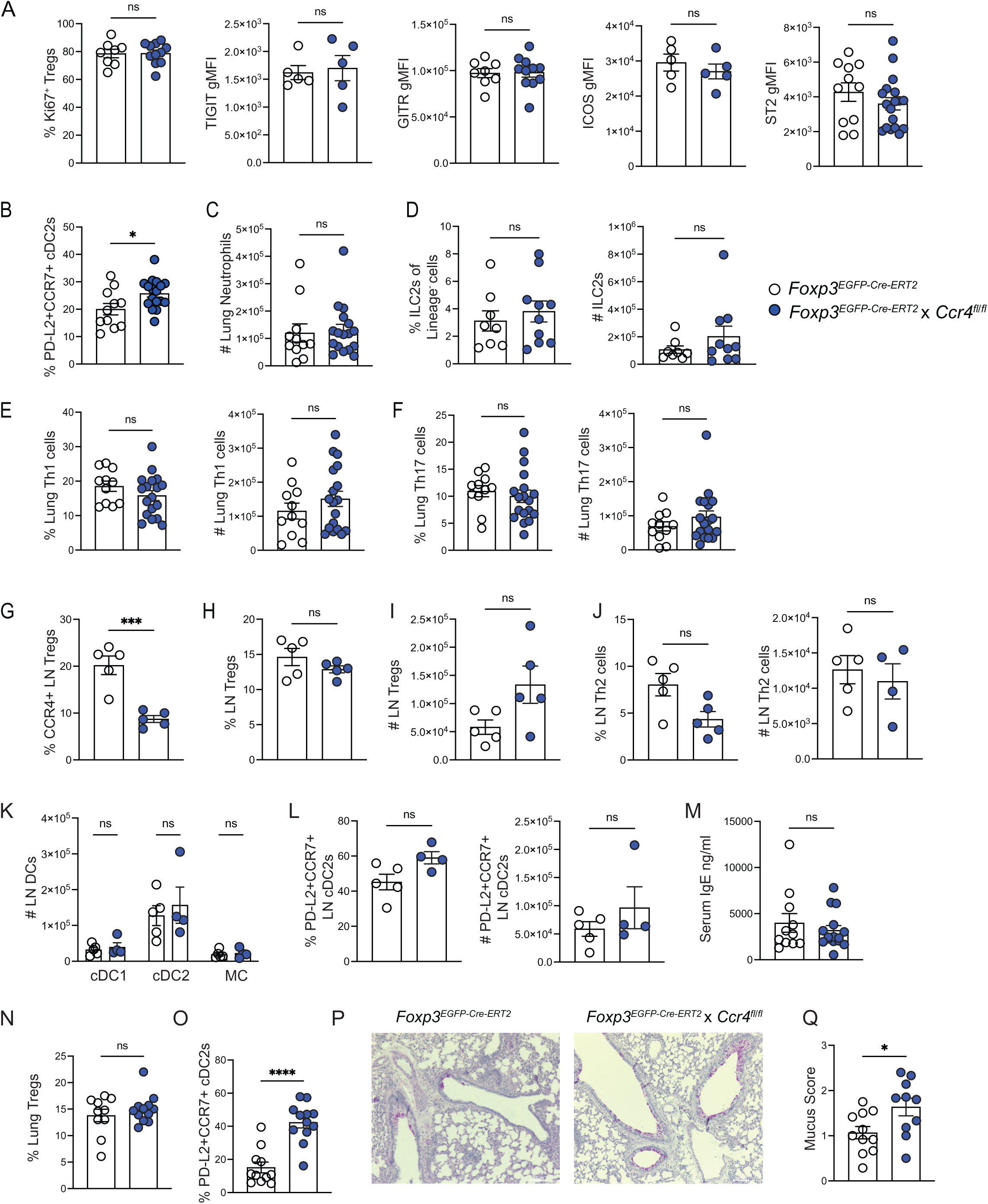
Immunophenotyping the allergic immune response upon deletion of CCR4 in Tregs. *Foxp3^EGFP-Cre-ERT2^* control mice and *Foxp3^EGFP-Cre-ERT2^* x *Ccr4^fl/fl^* mice were sensitized with 10 µg i.n. HDM followed by daily challenges of 10 µg i.n. HDM on days 7–11, 1 mg daily doses of tamoxifen via oral gavage on days 4-8, then injected with anti-CD45 antibody i.v. 3 minutes prior to tissue harvest on day 14. **A**. % Ki67^+^ and gMFI for TIGIT, GITR, ICOS, and ST2 in lung Tregs from indicated groups. **B**. Percentage PD-L2^+^CCR7^+^ cDC2s of total lung cDC2s. **C**. Total number of lung neutrophils. **D**. Percentage of lung ILC2s of lineage^-^cells and total number of lung ILC2s. **E**. Percentage and number of lung Th1 cells. **F**. Percentage and number of lung Th17 cells. **G**. Percent LN CCR4^+^Foxp3^+^ Tregs. **H**. Percentage LN Tregs. **I**. Number of LN Tregs. **J**. Percentage and number of LN Th2 cells. **K**. Number of total LN DC subsets. **L**. Percentage and total number of PD-L2^+^CCR7^+^ of LN cDC2s. **M**. Serum IgE levels. **N-Q**. *Foxp3^EGFP-Cre-ERT2^* control mice and *Foxp3^EGFP-Cre-ERT2^* x *Ccr4^fl/fl^* mice were administered 10 µg i.n HDM 3 times per week for 6 weeks, 1 mg daily doses of tamoxifen via oral gavage on days 4-8, 18-22 and 32-36 of HDM treatment, then injected with anti-CD45 antibody i.v. 3 minutes prior to tissue harvest on day 42. **N**. Percentage of lung Tregs of total CD4^+^ T cells from indicated groups. **O**. Percentage PD-L2^+^CCR7^+^ cDC2s of total lung cDC2s. **P**. Representative PAS-stained lung sections. **Q**. Mucus scores. Data are from three independent experiments with 11-18 mice pooled or data are representative of one experiment with *n*L=L5 mice per group from three independent experiments. For statistical analysis, a two-tailed *t* test was performed for parametric data, and a two-tailed Mann-Whitney *U* test was performed for nonparametric data. One-way ANOVA analysis with Holm-Sidak’s testing for multiple comparisons. *, p<0.05; ****, p<0.0001, ns, not significant.

**Supplementary Figure 5.**
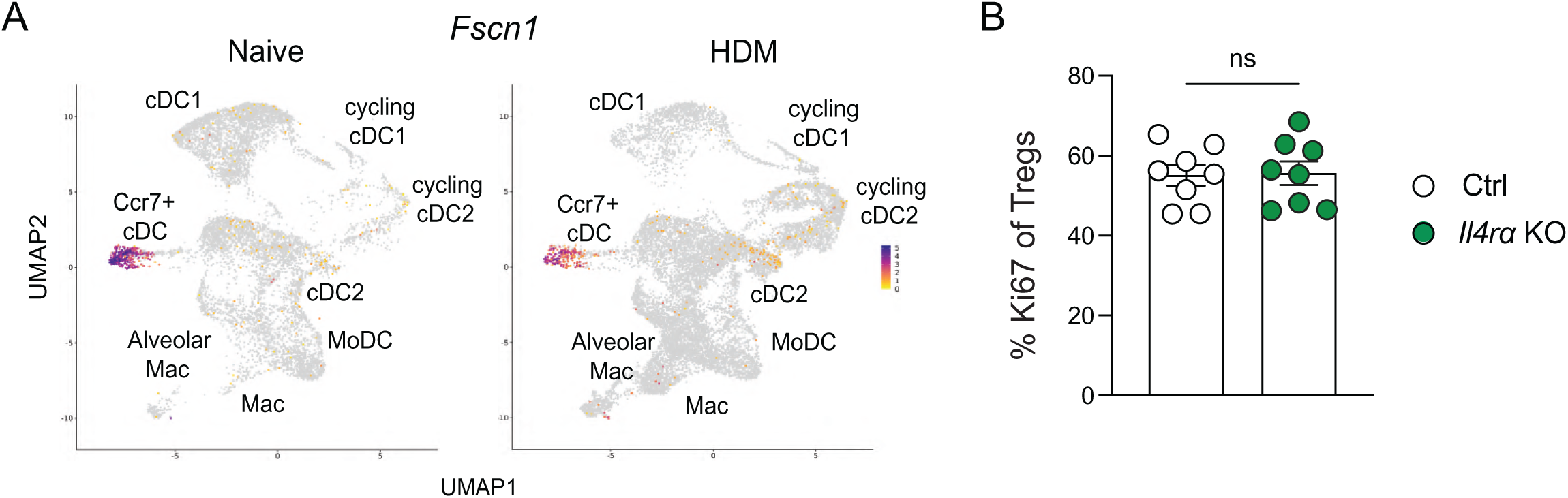
Expression of *Fscn1* in lung DCs and proliferate potential of Foxp3^+^ Tregs. **A**. UMAP of *Fcsn1* expression in lung DC clusters from nailJve and HDM-treated mice. **B**. Percent Ki67^+^ among lung Tregs from Ctrl:Il4ra KO mixed bone marrow chimera mice on day 14 of HDM treatment protocol.

